# A unique subset of pericystic endothelium associates with aberrant microvascular remodelling and impaired blood perfusion early in polycystic kidney disease

**DOI:** 10.1101/2024.03.03.583132

**Authors:** Daniyal J Jafree, Charith Perera, Mary Ball, Daniele Tolomeo, Gideon Pomeranz, Laura Wilson, Benjamin Davis, William J Mason, Eva Maria Funk, Maria Kolatsi-Joannou, Radu Polschi, Saif Malik, Benjamin J Stewart, Karen L Price, Hannah Mitchell, Reza Motallebzadeh, Yoshiharu Muto, Robert Lees, Sarah Needham, Dale Moulding, Jennie C Chandler, Claire L Walsh, Adrian S Woolf, Paul J D Winyard, Peter J Scambler, René Hägerling, Menna R Clatworthy, Benjamin D Humphreys, Mark F Lythgoe, Simon Walker-Samuel, David A Long

**Affiliations:** Developmental Biology & Cancer Research & Teaching Department, UCL Great Ormond Street Institute of Child Health, UCL, London, UK; UCL Centre for Kidney and Bladder Health, University College London, London, UK; Specialised Foundation Programme in Research, NHS East of England, Cambridge, UK; UCL Centre for Advanced Biomedical Imaging, London UK; Central Laser Facility, Science and Technologies Facilities Council, UK Research and Innovation, Didcot, UK; Lymphovascular Medicine and Translational 3D-Histopathology Research Group, Charité Universitätsmedizin Berlin, Berlin, Germany; Berlin Institute of Health at Charité-Universitätsmedizin Berlin, BIH Center for Regenerative Therapies, Berlin, Germany; Molecular Immunity Unit, Department of Medicine, University of Cambridge, Cambridge, UK; Wellcome Sanger Institute, Wellcome Genome Campus, Hinxton, United Kingdom; Mathematical Sciences Research Centre, Queen’s University Belfast, Belfast, UK; Research Department of Surgical Biotechnology, Division of Surgery and Interventional Science, UCL, London, UK; UCL Institute of Immunity and Transplantation, UCL, London, UK; Division of Nephrology, Department of Medicine, Washington University in St. Louis, St. Louis, MO, USA; Department of Mechanical Engineering, University College London, London, UK; School of Biological Sciences, Faculty of Biology Medicine and Health, University of Manchester, Manchester, UK

## Abstract

Hallmarks of autosomal dominant polycystic kidney disease (ADPKD), the most common hereditary kidney anomaly, include expanding fluid-filled epithelial cysts, inflammation, and fibrosis. Despite previous work showing the potential of vascular-based therapies, renal microvascular alterations in ADPKD, and their timing, are poorly understood. Using single-cell transcriptomics of human kidney microvasculature, we identify a population of endothelial cells adjacent to cysts in ADPKD. This pericystic endothelium, distinguishable by its expression of osteopontin (SPP1), has a distinct molecular profile compared to the common endothelial cell injury signature in other kidney diseases. SPP1^+^ pericystic endothelium was also present in an orthologous mouse model of ADPKD before overt kidney functional decline. By interrogating geometric, topological and fractal properties from three-dimensional imaging of early ADPKD mouse kidneys, we show that pericystic endothelium associates with disorganisation and non-uniformity of the renal cortical microvasculature. Concurrently, we detected region-specific reductions in cortical blood flow within ADPKD murine kidneys using arterial spin labelling. We conclude that ADPKD kidneys contain a unique subset of endothelium manifesting with aberrant remodelling and impaired blood perfusion. Its detection, prior to renal functional decline, advocates the vasculature as a therapeutic target to modulate or preserve renal function in early ADPKD.

## INTRODUCTION

Autosomal dominant polycystic kidney disease (ADPKD) is the most common hereditary renal disease (1), and characterised by fluid-filled renal cysts which expand insidiously throughout the patient’s lifespan. This is associated with inflammation and fibrosis, resulting in replacement of the normal parenchyma of the kidney by cystic tissue, frequently resulting in kidney failure. Most cases of ADPKD are driven by mutations in *PKD1* or *PKD2* (2), encoding polycystin proteins that are enriched in primary cilia (3). Given that polycystins primarily localise to renal tubular epithelium (4, 5) and cysts are epithelial in origin, our pathophysiological understanding of ADPKD is focused towards cellular and molecular alterations to epithelial cells. Mutations in *PKD1* or *PKD2* alter renal tubular mechanosensation and disrupt intracellular calcium homeostasis (5, 6). This results in increased levels of cyclic adenosine monophosphate (cAMP) (7, 8), dysregulated epithelial cell proliferation (9, 10), ion secretion (11) and cyst formation. Current drugs for ADPKD are directed towards the kidney epithelium. The clinically licensed drug, tolvaptan, targets tubular arginine vasopressin-mediated cAMP (12), whereas somatostatin mediates inhibition of epithelial chloride transport (13). Other interventions within the translational pipeline target epithelial cell metabolism (14). Despite these pharmacological approaches, many patients still progress to kidney failure (15), and so alternative targets that could complement available epithelial-targeted drugs are needed.

Mounting evidence suggests that non-epithelial cell types in the microenvironment surrounding cysts are important players in ADPKD (16–19). Of these, the microvasculature may represent a potential therapeutic target (20). The blood microvasculature, composed of endothelial cells (EC), performs canonical functions, including tissue oxygenation and nutrient delivery. However, in the kidney, the microvasculature is molecularly and structurally specialised to meet the diverse physiological demands of the organ, including ultrafiltration and solute reabsorption (21). There are several lines of evidence implicating the vasculature in ADPKD. Patients with ADPKD have vascular malformations, including aneurysms, in different organs (22). Targeted deletion of polycystins in the endothelium of embryonic mice results in abnormal vessel migration and patterning. By contrast, deleting endothelial polycystins in adult mice alters systemic vascular resistance and blood pressure (23, 24). These findings indicate that polycystin loss of function has specific consequences on ECs and the systemic vasculature. How the specialised vasculature of the kidney is affected by ADPKD remains elusive. In human kidneys with end-stage ADPKD, angiography, immunostaining and corrosion casting have all demonstrated the presence of extensive, tortuous and disorganised blood vascular networks within the tissue microenvironment surrounding cysts (25, 26), findings which have been mirrored in rodent models (27–29). Such kidney microvascular alterations could occur in late disease from compression by expanding, fluid-filled cysts. An alternative hypothesis is that vascular alterations in ADPKD occur early, alongside cyst growth, and modify disease severity (20, 30). For example, peritubular capillaries density increases early in mouse kidneys with rapidly progressive ADPKD (29). However, the nature of these structural alterations is unclear, and the molecular and functional consequences of ADPKD on the kidney microvascular vasculature are completely unknown. The timing of such changes in relation to progression of cystic disease is also unclear.

To assess the molecular profile of the microvasculature in cystic kidneys, we harnessed single-cell transcriptomics of human ADPKD tissues, paired with three-dimensional (3D) confocal microscopy. We sought to ascertain whether molecular changes were mirrored in kidneys from mice which carry a hypomorphic *Pkd1* p.R3277C point mutation (RC) identified in patients (31, 32) in homozygosity (*Pkd1^RC/RC^*). In parallel, 3D structural information on vascular architecture was obtained using a novel geometric, topological and fractal analysis framework applied to 3D confocal microscopy of *Pkd1^RC/RC^* kidneys. Functional information on kidney vascular perfusion was ascertained using arterial spin labelling (ASL) magnetic resonance imaging (MRI). By using this multimodal approach, we identified a subset of the blood microvasculature unique to ADPKD. This endothelial population has a distinct molecular profile suggesting metabolic and angiogenic abnormalities, and maps to the region adjacent to cysts in diseased humans and mice. We therefore describe it as pericystic endothelium. Within the cortex of *Pkd1^RC/RC^*mice, pericystic endothelium associates with aberrant patterning, featuring non-uniformity and disorganisation, and impaired blood perfusion. Critically, these changes occurred prior to inexorable decline in renal function of the mouse model, advocating the vasculature as a therapeutic target in early ADPKD.

## METHODS

### Ethics statement

Ethical approval to obtain ADPKD tissue samples for 3D imaging was granted by the UK National Research Ethics Committee (21/WA/0388) and approved by The Royal Free London NHS Foundation Trust-UCL Biobank Ethical Review Committee (NC.2018.007, B-ERC-RF). Mouse experiments were carried out according to a UK Home Office project license (PPL: PP1776587) and were compliant with the UK Animals (Scientific Procedures) Act 1986.

### Analysis of single-cell and single-nucleus RNA sequencing data

#### Sample acquisition and pre-processing

We generated a transcriptomic atlas of the renal microvasculature using previously published single-cell RNA sequencing (scRNA-seq) data of healthy and diseased human kidneys (33). Healthy samples constituted tissues from donor kidneys which were not suitable for transplantation, non-rejection biopsies from kidney transplants or non-tumorous regions from tumour nephrectomies. Diseased human kidneys included those from individuals with chronic kidney disease and hypertension with or without diabetes, and transplants with chronic alloimmune rejection, chronic pyelonephritis or focal segmental glomerulosclerosis. Independently, we used the markers from this atlas to re-annotate ECs from previously published single-nucleus RNA sequencing (snRNA-seq) dataset of healthy or ADPKD tissues (34) acquired from the Gene Expression Omnibus database (GSE185948). Analyses of these two datasets were performed separately in R using Seurat (35), and the code for these analyses are provided in the following GitHub repository: https://github.com/daniyal-jafree1995/PKD-endothelium. Raw count matrices from each dataset were isolated from clusters annotated as ECs, verified by the co-expression of blood EC-enriched genes: *cadherin* (*CDH*)*5* and *vascular endothelial growth factor receptor* (*VEGFR*)*1*. Thereafter, the raw counts were independently log normalised, scaled by expression of all genes detected and subject to principal component analysis. The number of principle components used for downstream analysis was determined by the point where principal components cumulatively contributed to 90% of the standard deviation. Integration of cells from each donor was performed using the Harmony package (36). Nearest-neighbour graph construction, unsupervised clustering and uniform manifold approximation and projection (UMAP) were performed using the *FindNeighbors*, *FindClusters* and *RunUMAP* functions, respectively. Any cell clusters recognised as non-blood ECs or low quality and non-specific were removed, and the above procedure was iterated until only blood ECs remained.

#### Cell type annotation, differential abundance and unbiased cross-dataset comparison

For scRNA-seq and snRNA-seq data, lists of differentially expressed genes (DEG) discriminating between blood EC clusters were generated. These lists were manually inspected and compared with previously published datasets identifying specialised subsets of vasculature within the mouse (37, 38) and human (39) kidney to annotate specialised renal blood EC subtypes. To assess the comparative abundance of cell types within snRNA-seq data, we performed differential abundance testing using miloR (40), visualising differences in log-fold change abundance of each cell type between healthy kidneys and those with ADPKD using a beeswarm plot. To compare cell type annotations between scRNA-seq and snRNA-seq data, the SingleCellNet (41) tool was used, setting scRNA-seq annotations as ‘reference’ and snRNA-seq annotations as ‘query’, and visually compared using a heatmap.

#### Computing differentially expressed genes, gene ontology and marker selection

DEGs between distinct blood vascular subsets, or between healthy and ADPKD kidneys, were computed using the *FindAllMarkers,* considering genes with an adjusted *p* value < 0.05 or average log-fold change (log_2_FC) value greater than 0.25. DEGs between healthy and ADPKD kidneys for each blood EC subset were visualised using the EnhancedVolcano package. Gene ontology (GO) analysis was performed using the PANTHER webtool for gene classification (42) as previously described (33). Marker genes were selected based on the following criteria: i) an adjusted *p* value > 0.05 from a Wilcoxon rank sum test; ii) a log_2_FC value greater than 1; iii) within the top ten DEGs when lists of genes were ordered by log_2_FC (iv) non-secreted proteins to be identifiable by subsequent immunofluorescence experiments; (v) specificity of the transcript for vascular ECs by visual inspection of feature plots and (vi) consistent expression across the same specialised EC subtype across three or more of the datasets used to generate the scRNA-seq atlas, examined using a dot plot.

### Mouse *in vivo* experiments and imaging

#### Husbandry

Animal experiments were performed according to the ARRIVE guidelines for reporting animals in research (43). To mimic the slow temporal progression of human ADPKD (31), we utilised mice carrying a p.R3277C (RC) hypomorphic allele of *Pkd1* (32), maintained on a C57BL/6 background for at least five generations. Mice carrying the RC allele in heterozygosity (*Pkd1^+/RC^*) were mated, and the progeny genotyped as previously described (32) to generate wildtype (*Pkd1^+/+^*) or homozygous (*Pkd1^RC/RC^*) mice. Mice were maintained up to 12 months of age and were sacrificed using terminal anaesthesia and death confirmed using cervical dislocation.

#### Surrogate markers of renal function

Kidney to body weight ratio and blood urine nitrogen (BUN) were utilised as surrogate markers for murine renal function and were analysed at multiple timepoints up to 12 months of age. Body weight was measured prior to sacrifice and weight of both kidneys was determined after sacrifice and harvesting of organs by midline laparotomy. BUN assays were performed using a commercially available kit (BioAssay Systems, Hayward, CA) as per manufacturer’s instructions.

#### Magnetic resonance imaging configuration

MRI protocols were applied to quantify kidney structure and renal blood flow (RBF). Prior to MRI, mice underwent anaesthetic induction using 4% isoflurane in 0.8 L/minute medical air and 0.2 L/minute O_2_. Following induction and weighing, mice were constrained into the MRI cradle in a supine position using electric tape. Anaesthesia was maintained during the acquisition by reducing isoflurane concentration to 2% in 0.4 L/minute medical air and 0.1 L/minute O_2_, with a scavenger pump inside the magnet bore to prevent isoflurane build-up. Body temperature was maintained at 37 ± 0.5°C using heated water tubing and monitoring using a rectal probe and breathing rate was maintained between 90-100 repetitions per minute and measured using a respiration pad (SA Instruments). Images were acquired on a 9.4 T Bruker imaging system (BioSpec 94/20 USR) with a horizontal bore and 440 mT/m gradient set with an outer/inner diameter of 205 mm/116 mm (BioSpec B-GA 12S2), 86 mm volume coil and a 2×2 four-channel array, 1H-receive-only mouse cardiac surface coil (RAPID Biomedical GmbH) for the transmission and the reception of the RBF signal, respectively.

#### Anatomical imaging using T_2_-weighted MRI

To perform *in vivo* anatomical imaging of the kidneys of *Pkd1^+/+^* or *Pkd1^RC/RC^* mice, respiratory-triggered, fat-saturated T_2_-weighted sequences were acquired with T_2_ TurboRARE sequence (fast-spin echo, Paravision v6.0.1; field of view = 20 mm × 30 mm; matrix size = 200 × 300; RARE factor 4, averages = 4, repetitions = 1; effective echo time, 15 ms; repetition time, 1,500 ms; slice thickness, 0.5 mm). This enabled us to differentiate cortex from medulla and facilitating positioning of subsequent RBF measurements.

#### Imaging of renal blood flow using arterial spin labelling

RBF was quantitatively mapped using a fat-saturated flow alternating inversion recovery (FAIR) pulsed ASL with an echo planar imaging (EPI) readout (field of view, 20 mm x 30 mm; matrix, 100 x 150, repetitions = 10; effective echo time, 16000 ms; repetition time, 20.15 ms; dummy scans, 2; imaging slice thickness, 2 mm; segments, 2; repetitions, 10). For quantification of RBF, an optimized inversion time of 2000 ms was used (44). Slice-selective inversion pulses were applied in the coronal plane matching the imaging slice with a bandwidth of 5200 Hz and thickness of 8 mm. Voxel-wise T_1_ and M_0_ values were also acquired by obtaining EPI data at multiple TI values. For this acquisition, only non-selective inversion was applied using identical parameters as for ASL. Multiple TI values were arrayed within a single sequence: TI = [30 50 100 150 200 300 500 1000 1500 2500 5000 8000 ms].

#### Analysis of MRI data

Regions of interest (ROI) were drawn manually on control (M_C_, non-selective) images to cover the renal cortex or medulla of each kidney, guided by T_2_-weighted imaging. T_1_ and M_0_ values for each voxel within the ROIs were determined using the T_1_ map EPI images. The averaged, multi-TI, non-selective data (M_C_) were fit to a simple inversion recovery model: M_C_ = M_0_ (1 – 2^−TI/T1^). Control (non-selective) and labelled (slice-selective) ASL images were averaged across the ten repetitions. For each ASL image pair, for each voxel within the ROI, the mean M_C_ was subtracted from the mean slice-selective value to provide the perfusion-weighted signal, ΔM. Voxel-wise values of renal perfusion within the ROI were then quantified from the ASL data using a previously described kinetic model (45, 46) which incorporates the averaged T_1_, M_0_, and ΔM values. The mean average RBF value from each mouse was reported, across cortex and medulla. T_2_-weighted imaging was used to guide manual selection of ROIs within the cortex to assess RBF within regions adjacent to macroscopically overt cysts.

### 3D imaging and analysis of the kidney and its blood microvasculature

#### Harvesting and storage of kidney tissues

For validation of scRNA-seq data, healthy human adult kidney tissue was obtained from three deceased organ donors who were not for transplantation (47), retrieved by a UK National Organ Retrieval Services team. For validation of snRNA-seq data, tissue was acquired from two individuals with ADPKD derived by the transplant surgical team. Once explanted, the tissues were pseudo-anonymised and incubated overnight in Belzer University of Washington Cold Storage Solution (Bridge to Life Europe, London, UK) at 4°C, before random subsampling into ∼2 mm^3^ pieces. Intact mouse kidney at 3 months of age in *Pkd1^+/+^* or *Pkd1^RC/RC^*mice was collected after MRI and kidney weight assessed after sacrifice. All kidney tissues were incubated in 4% (w/v) paraformaldehyde (PFA, Sigma Aldrich), made up in 1 X phosphate buffered saline (PBS), at 4°C overnight. All tissues were then washed and stored in 1 X PBS with 0.02% (w/v) sodium azide.

#### Immunofluorescence of healthy human kidney sections for marker validation

Human kidney donor tissues were dehydrated in ethanol, embedded in paraffin and cut into 5 μm sections utilising a rotating blade microtome. Antigen retrieval was performed using Tris-EDTA buffer and endogenous peroxidase quenched using 1.6% hydrogen peroxide. Slides were blocked using CAS-Block for 30 minutes before incubation overnight with the following primary antibodies: mouse monoclonal anti-platelet and endothelial cell adhesion molecule 1 (PECAM1/CD31 Aligent, GA610, clone: JC70A, 1:50) rabbit polyclonal anti-ssu-2 homolog (SSUH2, Atlas Antibodies, HPA049777, 1:100), rabbit polyclonal anti-fibulin 2 (FBLN2, ThermoFisher Scientific, PA5-51665, 1:200), rabbit polyclonal anti-glycoprotein M6A (GPM6A, Proteintech, 15044-1-AP, 1:100). AlexaFluor-conjugated secondary antibodies (ThermoFisher Scientific) used were for indirect immunolabelling at a concentration of 1:200, including donkey-anti mouse 546 and donkey anti-rabbit 546. Stained slides were mounted for imaging by Kohler illumination on a Zeiss Axioplan 2 scope.

#### Wholemount immunofluorescence and optical clearing

Human ADPKD kidney tissue samples were randomly selected from 2 mm^3^ pieces, with two samples used per patient. Three-month-old *Pkd1^+/+^*or *Pkd1^RC/RC^* mouse kidneys were manually cut into ∼ 500 μm^3^ slices in axial cross-section. The tissues were then subject to a wholemount immunostaining protocol (48) formulated by adaptations to iDISCO (49) and vDISCO (50) protocols. Unless otherwise stated, the following steps were performed at room temperature, all agitated on a flat shaker with incubation volumes of 1 mL, and reagents purchased from Sigma Aldrich. Kidney tissues were dehydrated in increasing concentrations of methanol (50, 70%) in double distilled (dd)H_2_O for 30 minutes per step, before bleaching in absolute methanol with 5% (v/v) of 30% hydrogen peroxide solution overnight at 4°C. The next day, tissues were rehydrated in the methanol series, followed by incubation in 1 x PBS for 30 minutes. Kidneys were permeabilised and blocked with antibody solution (0.09g trans-1-acetyl-4-hydroxy-L-proline, 0.033g methyl-β-cyclodextrin, 0.02450g sodium azide and 245 μL Triton X100 per 50 mL of 1 X PBS) with 6% donkey serum and 10% dimethyl sulfoxide (DMSO) overnight at 37°C, before incubation in antibody solution with 3% donkey serum and 5% DMSO with primary antibodies for three days at 37°C. Primary antibodies and labels used for wholemount staining in humans included mouse monoclonal anti-CD31 (1:50) and rabbit polyclonal anti-osteopontin (SPP1, Abcam, ab8448, 1:100). Rabbit polyclonal anti-osteopontin SPP1, rat monoclonal anti-CD31 (BD Biosciences, 557355, clone: MEC 13.3, 1:50), rat monoclonal anti-endomucin (EMCN, Santa-Cruz, sc-65495, clone: V.7C7, 1:50) and Dylight 649-conjugated *Lycopersicon esculentum* (Tomato) lectin (Vector Laboratories, DL-1178-1, 1:100) were used on mouse kidney slices. All tissues were then washed in wash solution (0.2 mL of 10mg/ml heparin stock, 0.4 mL Tween-20 and 0.01g sodium azide per 200mL of 1 x PBS) for four times for one hour per step, before incubation in antibody solution with 3% donkey serum and 5% DMSO with AlexaFluor secondary antibodies overnight at 37°C. All tissues were further washed in wash solution four times for one hour per step before dehydration in 50%, 70% and 100% methanol. Optical clearing was performed as previously described (48) using BABB (a 1:2 solution of benzyl alcohol and benzyl benzoate). Immunostained and dehydrated kidney tissues were equilibrated in a 1:1 solution of methanol:BABB, before incubation in BABB until full optical clearance.

#### Confocal microscopy

To capture high-resolution 3D images of the blood microvasculature, human ADPKD kidney tissue samples or 3-month-old *Pkd1^+/+^*or *Pkd1^RC/RC^* mouse kidney slices were placed between a large coverslip and cover glass, supported by a O-Ring (Polymax Ltd, Bordon, UK) made from BABB-resistant rubber, as described previously (48). Confocal images were acquired on an LSM880 upright confocal microscope (Carl Zeiss Ltd.), with 10×/NA 0.5 W-Plan Apochromat water dipping objective (working distance; WD = 3,700 μm). Using 488, 561 or 633 nm lasers for excitation, pericystic vasculature was imaged in three or more randomly selected non-overlapping ROI. In mouse samples, cortex and medulla were imaged separately, structurally discriminated by autofluorescent signal using the 488 nm laser. Gallium arsenide phosphide (GaAsP) internal and external detectors were used for high sensitivity. Image *z*-stacks were taken through up to 2 mm of tissue and exported as CZI files.

#### Lightsheet fluorescence microscopy

Images of cyst volume and distribution in 3-month-old *Pkd1^RC/RC^*mouse kidney slices were acquired using an Intelligent Imaging Innovations (3i) Cleared Tissue Light Sheet (CTLS) microscope. BABB-cleared samples were immersed in ethyl trans-cinnamate (ECi; 99%) for imaging. The glass imaging chamber was filled with ECi with the sample glued to a custom 3D-printed spoon using ultraviolet-activated glue (UV bonding Ltd.). The spoon was magnetically fixed to the stage and the sample immersed in the glass chamber with ECi. Illumination was provided by two 5x/0.14 NA air objectives (5X Mitutoyo Plan Apo Infinity Corrected Long WD Objective, #46-143, Edmund Optics) either side of the glass imaging chamber and the laser beam was swept perpendicularly to the detection objective to create a lightsheet from either side. A spatial light modulator was used to adjust for chromatic aberrations, as well as providing as many angles for the light as possible to reduce shadowing. For detection, a 1.0x/0.25 NA air objective (PlanNeoFluar Z 1.0x/0.25, Carl Zeiss Ltd.) was used, followed by a zoom module (Axio Zoom.V16, Carl Zeiss Ltd.) to change the lateral pixel dimensions on the sCMOS camera, with a zoom of 6.5x resulting in a lateral pixel dimension of 1 μm. The lightsheet focus was swept horizontally across the sample over five equally-spaced positions and an image was reconstructed to keep only the best axially-focused information, with an axial resolution of 7 μm. A 488 nm laser was used to image tissue structure. A quad-band emission filter was placed before the camera (Semrock FF01-446/523/600/677-25). Images were taken and exported as .sldy files.

#### Segmentation and analysis of cysts within mouse kidneys

Lightsheet images of 3-month-old *Pkd1^RC/RC^* mouse kidney slices were imported as .sldy files into FIJI (v2.1.0) (51). A 3D gaussian blur was applied to smooth the image stack before the Li thresholding was used to generate a binarized mask of the kidney. The mask was duplicated and on one duplicate, the Fill Holes function was used within the MorphoLibJ toolkit (52) to generate a filled mask of the kidney parenchyma. The original binarized mask, with cysts included, was subtracted from this image to generate a mask containing cysts only. Thereafter, the binarized cysts and the filled binarized kidney were saved as TIFF stacks and exported to IMARIS (Bitplane, v8.2). Within IMARIS, the Isosurface Rendering tool was used to semi-automatically segment the cysts and kidney volume as previously described (53), before manual inspection and removal of rendered structures corresponding to the renal pelvis and arterial vasculature. Total cyst number, cyst volume and kidney volume were then derived from the Statistics toolkit.

#### Segmentation and geometric analysis of kidney blood vasculature

Confocal microscopy stacks were imported as CZI images into FIJI. The rolling-ball algorithm was used to subtract background from the images before the image was re-scaled as isometric. The greyscaleClosingSphere function was applied within the CLIJ2 toolkit (54), before the Tubeness plugin was used to enhance filamentous structures of the vasculature. Thereafter, the 3D Simple Segmentation plugin (55) was leveraged to generate binarized masks of the vasculature in 3D. The resulting images were used for analysis of vascular branching metrics or global measures of vascular patterning, as described below. To derive vascular branching metrics from binarized masks, we utilised the open-source VesselVio (v1.1) application (56). Analysis was performed as per instructions (https://github.com/JacobBumgarner/VesselVio) in order to quantify geometric properties of the vascular network, including the length and radius of each vessel branch. We also derived the total number of segments within the vascular network and divided this by the total volume of the kidney volume imaged. Here, kidney tissue volume was quantified by generating a binarized mask of tissue autofluorescence using the method described above, before using the 3D Object Counter plugin in FIJI for quantification. Multiple representative ROIs were analysed per kidney, imaging cortex and medulla separately.

#### 3D fractal and topological analysis of the kidney blood vasculature

From binarized image stacks, we also derived global measures of patterning of the vasculature using fractal analysis and topological data analysis approaches. Briefly, fractal and connectivity dimensions were calculated as described previously (57). These parameters give measures of the complexity or “roughness” of an object or pattern with higher values denoting a more complex and connected vasculature structures. Lacunarity is a fractal based measure of the distribution of gaps within the vasculature network (58), with higher values suggesting heterogeneity due to the presence of voids or spatial clustering. For topological data analysis approaches, 3D skeletonized vasculature networks were first extracted from binary vascular image stacks using ImageJ FIJI skeletonize3d plugin. Next, the persistence diagram was calculated from branch starting and end points using the python gudhi package (version 3.9.0) from which parameters of interest were calculated as previously described (59). the first Betti number - representing the number of vascular loops in the dataset, persistence entropy - a measure of the diversity or variability of the persistence values where a higher value suggests the presence of a wider variety of topological features across different scales, and the average lifetime of topological features across the dataset. Values were calculated independently for each ROI.

#### Quantification of vascular osteopontin expression

We used FIJI to assess the fluorescence intensity of SPP1 expression in the cortical vasculature of 3-month-old *Pkd1^RC/RC^* mouse kidney slices and wildtype controls, which had been wholemount immunolabelled for EMCN and SPP1 and cleared in BABB before 3D confocal microscopy. A maximum intensity project was generated from each region of interest within the kidney cortex. We used the Measure tool to sample across the whole kidney tissue on average, and thereafter, randomly sampled a minimum of ten EMCN^+^ vascular branches per ROI. We derived the mean SPP1 fluorescence intensity such that 30-50 individual vessels per mouse were analysed.

#### Sample size and statistics

Based on our prior experiments using this mouse model of PKD (29), 5-8 mice have been sufficient to assess differences between BUN and kidney to body weight ratio. Due to its sensitivity (53) and based on empirical assessment from a small cohort of 3 mice per group, we required only four mice per group for sufficient statistical power of 3D vascular analysis. For ASL measurements, we performed pilot experiments of 3 mice per group to determine that 4-6 mice would be required to demonstrate the differences observed in RBF. Apart from single-cell transcriptomic analysis described above, all statistics were performed in Prism (v9.5, GraphPad). Continuous data were assessed for normality using Shapiro-Wilk tests and Brown-Forsythe tests were used to assess equality of variance. We used unpaired Student’s *t* tests to compare *Pkd1^RC/RC^* and *Pkd1^+/+^*mice for each parameter, with one-way ANOVA and Tukey’s pairwise multiple comparisons where more than one timepoint, or region of kidney, was compared. Statistically significant differences were deemed to be *p ≤* 0.05 and are presented with mean differences, standard error of the mean (SEM) and 95% confidence intervals (CI) for each difference.

## RESULTS

### Single cell transcriptomic meta-analysis of human kidney vasculature

Before defining how the vasculature is altered in ADPKD, a reference of the transcriptional signatures of each segment of the human kidney vasculature was required. The kidney is perfused by a hierarchically arranged system of arterial vessels and arterioles. In the cortex, ultrafiltration is performed by glomerular vasculature and solute reabsorption by peritubular capillaries, whereas in the medulla, urinary concentration occurs by solute exchange between vasa recta and specialised tubular epithelium (**Fig.1A**). To transcriptionally resolve all these subsets and define a census of EC molecular markers that could be applied to any dataset, we performed a meta-analysis of existing scRNA-seq data featuring 41,546 blood EC, defined by co-expression of pan-endothelial markers *CDH5* and *VEGFR1,* pooled across a total of 64 human kidneys (**Fig.S1A**) (47). These included 25,072 ECs from healthy kidneys and 16,474 ECs from kidneys with different aetiologies of chronic kidney disease (60) or chronic transplant rejection (47, 61).

**Fig. 1.**
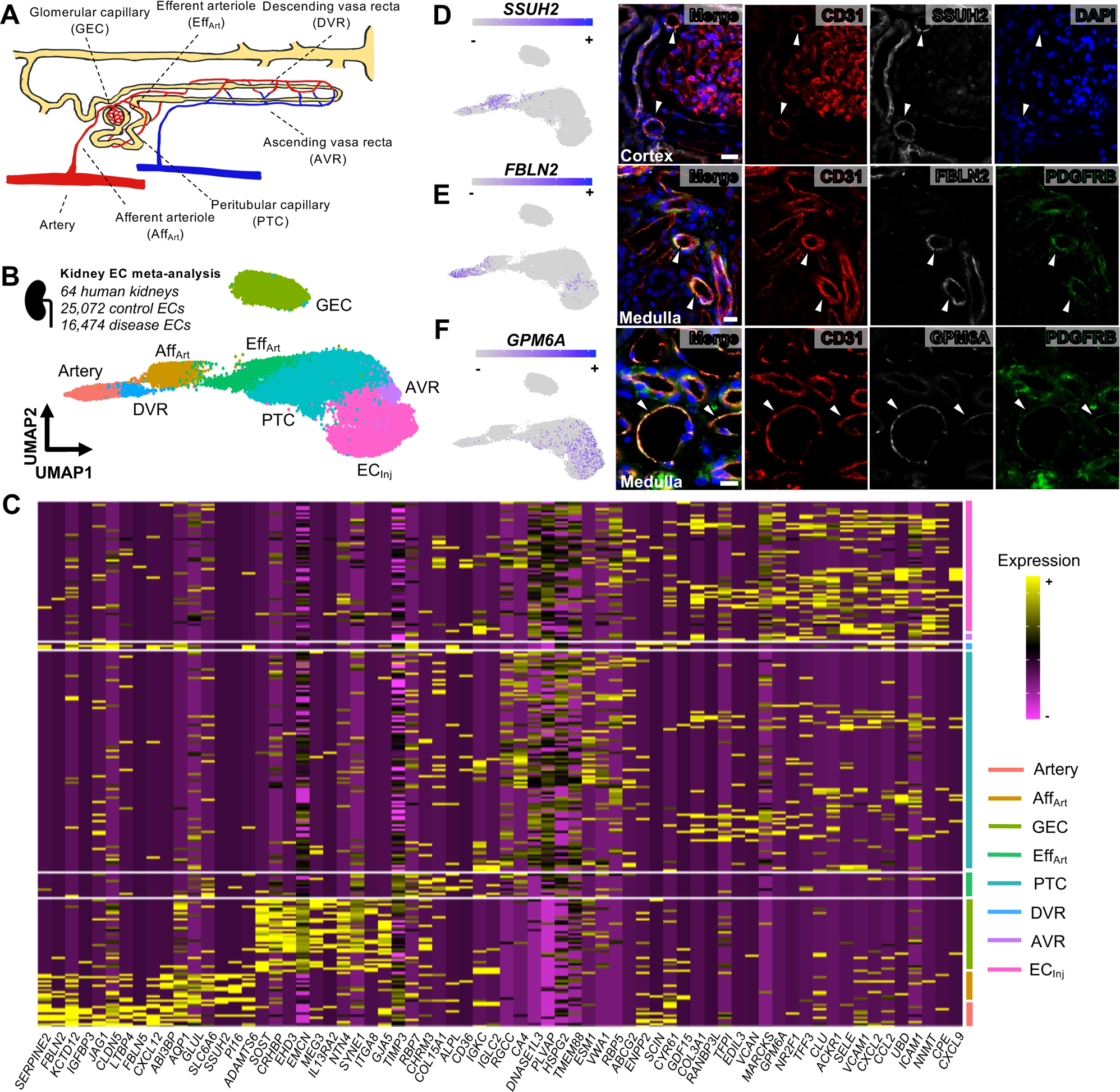
A single-cell RNA sequencing census of blood vasculature in the human kidney. **(A)** Schematic demonstrating the specialised blood vascular subsets within the kidney including arteries, afferent (Aff_Art_) and efferent (Aff_Eff_) arterioles, glomerular capillaries (GEC), peritubular capillaries (PTC) and ascending (AVR) and descending vasa recta (DVR). The tubular nephron is shown in yellow for spatial orientation. **(B)** Uniform manifold approximation and projection (UMAP) of 41,546 blood endothelial cells (EC) sampled across a total of 64 human kidneys from publicly available single-cell RNA sequencing (scRNA-seq) data. Seven annotated clusters correspond to the annotations in (**A**). A further cluster, deemed to be inflammation-activated ECs (EC_Inj_) was resolved. **(C)** Heatmap showing ten differentially expressed genes (DEG) for each cluster in the scRNA-seq data. Yellow indicates high expression and purple indicates low expression. Territories corresponding to each cluster are shown on the right axis. **(D)** Ssu-2 homolog (SSUH2) is a marker of the afferent arterioles (log_2_FC = 2.19, adjusted *p* < 0.0001), shown by UMAP on the left, validated by confocal imaging of immunostained human kidney paraffin sections stained for CD31 and DAPI on the right. Expression of SSUH2 is shown in CD31-expressing vessels adjacent to the glomerulus. Images are representative of five slides analysed across *n* = 3 human kidneys. Scale bar = 50 μm. **(E)** Fibulin 2 (FBLN2) is a marker of the descending vasa recta (log_2_FC = 1.55, adjusted *p* < 0.0001), shown by UMAP on the left, validated by confocal imaging of immunostained human kidney paraffin sections stained for CD31 and the pericyte marker, PDGFRβ, on the right. Expression of FBLN is shown in CD31-expressing vessels within the medulla which have PDGFRβ^+^ cell coverage. Images are representative of five slides analysed across *n* = 3 human kidneys. Scale bar = 50 μm. **F**) Glycoprotein M6A (GPM6A) is a marker of the ascending vasa recta (log_2_FC = 1.24, adjusted *p* = 1.17 × 10^−68^), shown by UMAP on the left, validated by confocal imaging of immunostained human kidney paraffin sections stained for CD31 and the pericyte marker, PDGFRβ, on the right. Expression of FBLN is shown in CD31-expressing vessels within the medulla which do not have PDGFRβ^+^ cell coverage. Images are representative of five slides analysed across *n* = 3 human kidneys. Scale bar = 50 μm.

We transcriptionally resolved eight distinct clusters of blood ECs (**Fig.1B**). Computation of DEGs (**Fig.1C**) provided lists of molecular markers (**Table S1**) that were used to annotate each cluster, informed by previous studies molecularly characterising kidney endothelial subsets in mouse (37, 38) and human (39). Seven of the eight clusters corresponded to anatomically distinct segments of the kidney blood vasculature. Glomerular capillaries clustered separately from the remainder of ECs, expressing established glomerular capillary markers including EH domain containing 3 (*EHD3*) (37, 62, 63). Another EC cluster was enriched for the arterial elastic lamina extracellular matrix components *FLBN2* and *FLBN5* (64), alongside the arterial endothelium-enriched chemokine C-X-C motif chemokine ligand (*CXCL*)*12*, (65) consistent with an arterial identity. The remainder of the EC clusters did not have as clear transcriptional distinction, and no individual marker was completely specific for any given EC cluster. Afferent arterioles were classified based on previous studies that identified their ECs to express peptidase inhibitor 16 *(PI16)* (38) and solute carrier family (SLC) 6 member 6 (*SLC6A6*) (37). ECs expressing not only arterial markers such as *CXCL12*, but also plasmalemma vesicle associated protein (*PLVAP*)(39) and tissue inhibitor of matrix metalloproteinase 3 (*TIMP3*)(66), were classed as efferent arteriole. Peritubular capillaries also expressed *PLVAP,*(37, 38) whilst concurrently expressing carbonic anhydrase 4 (*CA4*) (67). With regard to the medullary segments of the vasculature, the descending vasa recta shared arterial markers such as *FBLN2* (39), whilst also expressing *SLC14A1* (38), whereas the ascending vasa recta were characterised based on the presence of insulin-like growth factor binding protein (*IGFBP*)7, with scant *IGFBP5* transcripts detected (37). We further observed a cluster that was unique to the 16,474 ECs from diseased kidneys and was enriched for transcripts associated with inflammatory-mediated activation of ECs such as inflammatory cell adhesion molecule 1 (*ICAM1*) and vascular cell adhesion molecule 1 (*VCAM1*) (68). As these *VCAM1^+^ ICAM1^+^* ECs were clustered together irrespective of the disease they came from, they likely represent a common transcriptional signature of kidney endothelial injury, and we therefore labelled them as ‘injured’ endothelial cells (EC_Inj_).

To validate our annotations, we assessed the expression of novel marker genes within selected EC populations that had not been previously reported in the literature and validated these at the protein level using immunofluorescence of tissue sections from human kidney. We selected three genes based on their high differential expression with log_2_FC value > 1, their specificity for ECs as opposed to other non-EC cell types within the kidney dataset (**Fig.S1B**) and their commonality across multiple datasets and donors in the meta-analysis (**Fig.S1C**). One of the top candidates identified in the afferent arteriolar cluster was Ssu-2 homolog (*SSUH2*, log_2_FC = 2.19, adjusted *p* < 0.0001), a transcriptional regulator associated with odontogenesis (69). Correspondingly, SSUH2 was detected in large diameter cortical CD31^+^ vessels adjacent to glomeruli of human kidneys (**Fig.1D**). Conversely, *FBLN2*, which is expressed by large arteries in our dataset, was also enriched in descending vasa recta ECs (log_2_FC = 1.55, adjusted *p* < 0.0001), whereas ascending vasa recta ECs were enriched for expression of glycoprotein M6A *(GPM6A,* log_2_FC = 1.24, adjusted *p* = 1.17 × 10^−68^). We validated these findings using immunofluorescence, utilising platelet derived growth factor receptor beta (PDGRFβ) staining to discriminate mural cells of CD31^+^ ascending vasa recta. Accordingly, FBLN2 was co-expressed by descending vasa recta (**Fig.1E**), whereas GPM6A was detected in CD31^+^ medullary vessels without PDGRFβ^+^ mural cell coverage; the ascending vasa recta (**Fig.1F**). Thus, by transcriptionally resolving blood EC subsets in the kidney, we have identified and validated novel molecular markers which may serve to reproducibly annotate other single-cell transcriptomic datasets.

### A unique blood vascular phenotype in pericystic regions of human end-stage ADPKD

Having molecularly defined the known subsets of blood ECs within the human kidney, we then applied these findings to re-annotate snRNA-seq data of the vasculature in human ADPKD. Data from a recent snRNA-seq study (34) were analysed, comprising five control kidneys (*n =* 806 ECs) and eight kidneys with ADPKD explanted after kidney failure (*n =* 2,974 ECs) totalling 3,780 EC nuclei (**Fig.S2A**). The nuclei resolved into eight transcriptionally distinct clusters (**Fig.2A**), with DEGs for each cluster shown in (**Table S2**). These clusters included all the subsets derived from our earlier single-cell transcriptome meta-analysis, including *FBLN2*^+^ arteries and descending vasa recta, a hybrid afferent / efferent arteriolar cluster enriched for *SSUH2*, and *GPM6A^+^* ascending vasa recta (**Fig.S2B**). Glomerular or arterial ECs, efferent and afferent arterioles and ascending vasa recta mapped were detected in both control and ADPKD kidneys. Conversely, both visual inspection of the UMAP by (**Fig.2B**), quantification of cell types in each dataset (**Fig.S2C**) and through differential abundance analysis (40) (**Fig.2C**), we detected an uneven distribution of other ECs types in ADPKD as compared with normal kidneys. Descending vasa recta and peritubular capillaries were predominantly detected in ADPKD kidneys, likely to be a combination of sampling bias, the relatively low number of ECs sampled in healthy kidneys and the depletion of peritubular capillary ECs by snRNA-seq (39). We resolved two further EC subsets enriched in ADPKD kidneys. Akin to our scRNA-seq meta-analysis which identified a common kidney endothelial injury signature, one cluster was enriched for markers of EC injury or adhesion, including *ICAM1* and *VCAM1*. However, the second subset did not map to any cluster detected in our earlier meta-analysis. To verify this in an unbiased fashion, we trained a Random Forest classifier (41) using the ECs annotations in the meta-analysis and used these to assess ADPKD EC subtypes captured in the snRNA-seq. This analysis is shown in a classification heatmap, where each bar represents a classification score indicating similarity between cell types (**Fig.2D**). We identified similarity between common EC subtypes in the ADPKD scRNAseq atlas as compared to the previous scRNAseq reference, including GECs, arterial ECs, afferent and efferent arteriolar ECs and ascending or descending vasa recta. Some transcripts were differentially expressed between classical kidney EC subsets in control and ADPKD tissues (**Fig.S2D**). Injured endothelial cells also shared transcriptionally similar signatures between the meta-analysis and the ADPKD data. The unknown subset of ADPKD-enriched ECs, however, were not transcriptionally specific for any kidney EC subset in health or diseased samples. Given the exclusive presence of these unknown ECs in ADPKD kidneys, and their distinct transcriptional profile from the common *ICAM1^+^ VCAM1^+^* EC injury signature, we referred to these unique cells as ‘pericystic’ endothelium (EC_Cyst_).

**Fig. 2.**
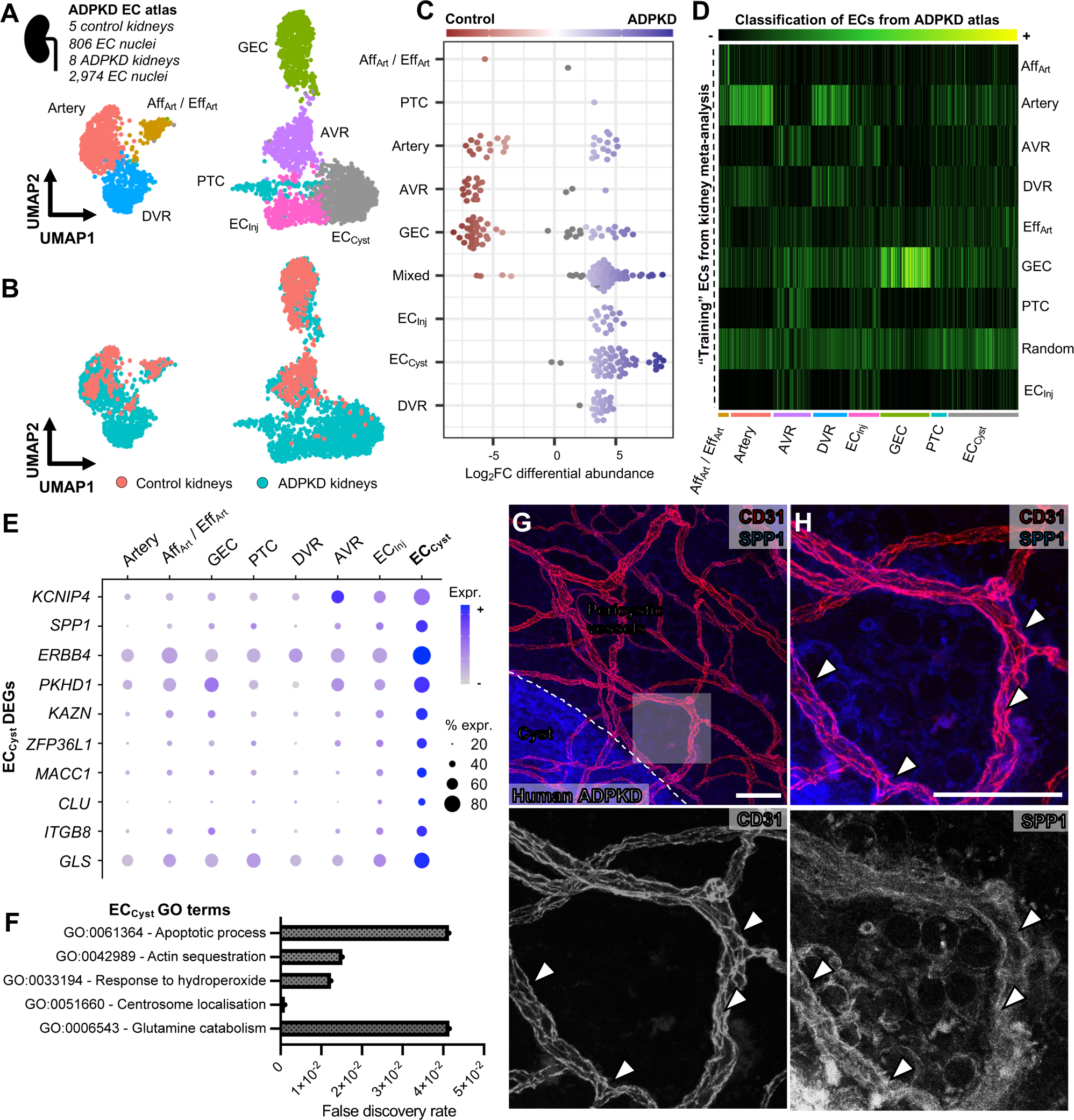
Annotation of single-nucleus RNA sequencing data of human ADPKD demonstrated a unique population of pericystic endothelium. **(A)** UMAP of 3,780 blood EC nuclei sampled across a total of 5 human control kidneys and 8 ADPKD explants from publicly available single-nucleus RNA sequencing (snRNA-seq) data. Eight transcriptionally distinct clusters were detected, including annotations shown in Fig.1 with a further unique cluster shown in grey. These were annotated as pericystic endothelium (EC_Cyst_). **(B)** The UMAP in (**A)** is grouped by sample as a visual demonstration of cell type abundance between conditions. **(C)** Differential abundance analysis using the Milo package, showing distributions of each cellular ‘neighbourhood’ within the samples. There is an overrepresentation of certain cell types in ADPKD, including injured endothelial cells, pericystic endothelium, and DVR. **(D)** Random forest classification using the SingleCellNet tool, with the reference being the scRNA-seq data shown in Fig.1, and the query dataset being the ADPKD snRNA-seq atlas of ECs. Each cell within the ADPKD atlas is assigned a transcriptional similarity score to known cell types within the scRNA-seq reference. The lightness of the green in the heatmap represents the score, and thus indicates transcriptional similarity to a given cell type. **(E)** Dot plot of top DEGs detected in pericystic endothelium, the size of the dot indicates the proportion of cells expressing the DEG, and the darkness of colour indicates the average level of expression within the cell cluster. **(F)** Gene ontology (GO) analysis of all DEGs within the pericystic endothelial cluster, with GO terms shown on the y axis and false discovery rate (FDR) shown on the x axis. **(G)** Representative confocal microscopy of *n* = 2 human ADPKD explants stained for CD31 and SPP1 before optical clearing. The dotted line demarcates SPP1-expressing cyst epithelium, and the transparent box indicates a higher-magnification region of interest shown in (**H**). Scale bar = 100 μm. **(H)** High magnification region of interest from (**G**) showing expression of SPP1 in the pericystic CD31^+^ vasculature. Regions of overlap are shown by white arrowheads. Scale bar = 100 μm.

To interrogate the molecular phenotype of pericystic endothelium more closely, we inspected the top DEGs for this cluster as compared with all other clusters in the ADPKD snRNA-seq dataset (**Fig.2E**). Several identified DEGs have plausible relationships to ADPKD pathogenesis and progression, with the top three candidates by adjusted *p-*value discussed below. Kv channel-interacting protein 4 (*KCNIP4)* was enriched in pericystic endothelium (log_2_FC = 0.71, adjusted *p* = 1.4 × 10^−70^); a potassium channel-interacting protein which responds to intracellular calcium levels (70), which are known to be impaired in *Pkd1*-deficient ECs (71). The inflammatory mediator (72) and renal disease biomarker (73–75), osteopontin (*SPP1*) was also enriched in pericystic endothelium (log_2_FC = 1.22, adjusted *p* = 6.26 × 10^−63^), as was Erb-B2 receptor tyrosine kinase 4 (*ERBB4,* log_2_FC = 0.83, adjusted *p* = 1.7 × 10^−58^), the expression of which modulates cyst growth in mice and human cyst epithelial cell lines (76, 77). Pericystic endothelium was also enriched for *PKHD1*, which encodes fibrocystin, interacts with *PKD1* (78) is associated with autosomal recessive PKD (log_2_FC = 0.96, adjusted *p* = 1.1 × 10^−56^), although *PKD1* itself was not within the list of DEGs. Hypoxia inducible factor (HIF)1A (log_2_FC = 0.48, adjusted *p* = 3.0 × 10^−13^) was also enriched in pericystic endothelium; noteworthy given previous reports of regional hypoxia in PKD driving HIF accumulation postulated to contribute to pericystic hypervascularity (79).

We then performed GO analysis of the DEGs identified in the pericystic endothelial subset (**Fig.2F**). Pericystic endothelium was enriched for pathways involved in glutamine catabolism (FC = 63.3, FDR = 4.2 × 10^−2^), apoptosis (FC = 63.3, FDR = 4.1 × 10^−2^) and GO terms relating to polarity, including centrosome localisation (FC = 38.0, FDR = 1.1 × 10^−3^) and actin sequestration (FC = 28.5, FDR = 1.5 × 10^−2^). Confirming the uniqueness of the pericystic endothelial phenotype to ADPKD, we found that the top DEGs enriched in pericystic endothelium were detected at scant levels, or were not expressed, in our scRNA-seq reference atlas of human healthy and diseased kidney microvascular ECs (**Fig.S2E**), with the exception of the RNA-binding protein butyrate response factor 1 (ZFP36L1), which is required for normal vascularisation (80). We further assessed genes expressed by pericystic endothelium at lower levels on average compared to other cell types within the snRNA-seq dataset (**Table S3**). Pericystic endothelium had significantly lower expression of vascular tyrosine kinases essential for normal vascular development and angiogenesis (20, 81) (**Fig.S2F**). These include members of the vascular endothelial growth factor receptor (VEGFR) family, *VEGFR1* (log_2_FC = −0.72, adjusted *p* = 4.63 × 10^−21^) and *VEGFR2* (log_2_FC = −0.53, adjusted *p* = 4.01 × 10^−10^), but not *VEGFR3* (adjusted *p* > 0.05) and tyrosine kinase with immunoglobulin-like and EGF-like domains *(TIE)2* (log_2_FC = −0.81, adjusted *p* = 1.03 × 10^−30^) and, to a lesser degree, *TIE1* (log_2_FC = −0.26, adjusted *p* = 1.53 × 10^−6^).

We then sought to validate the presence of pericystic endothelium *in vivo*. Fresh cystic kidney tissue pieces of ∼2 mm^3^ volume were obtained after total nephrectomy from two individuals with late-stage ADPKD, before applying wholemount immunolabelling for the blood vascular marker CD31, optical clearing and confocal 3D imaging (48, 53). SPP1 was selected as a candidate pericystic endothelial marker and labelled for, given that is has not previously known to be expressed by ECs in this context (73). Upon 3D imaging of optically cleared ADPKD tissue, we observed disorganised networks of CD31^+^ blood vasculature in close proximity to SPP1^+^ cyst-lining cells (**Fig.2G**). Upon higher magnification of pericystic regions, we observed co-expression of CD31 and SPP1 within the microvasculature immediately adjacent to cysts (**Fig.2H**). Thus, we identified a unique population of endothelium in the pericystic regions of human ADPKD which is transcriptionally and phenotypically distinct and unique to this disease. Our molecular analysis collectively suggests that pericystic endothelium can be recognised by its expression of SPP1, and possesses a transcriptome associated with abnormal metabolism and impaired angiogenesis within the microenvironment of ADPKD kidneys.

### Pericystic microvessels in early murine PKD are associated with aberrant vascular remodelling

In human ADPKD, kidneys tend only to be removed in advanced stages of the disease, precluding the analysis of earlier timepoints in the disease process. We therefore turned to a representative animal model which would enable us to test the hypothesis that abnormal vasculature was present at early disease stages, prior to inexorable decline in kidney function. *Pkd1^RC/RC^*mice mimic the slow temporal progression of human ADPKD as compared to other more rapidly progressive mouse models (31, 32). Histological analysis of *Pkd1^RC/RC^*mouse kidneys demonstrated the presence of epithelial cysts as early as 3 months of age (**Fig.3A**), with a corresponding increase in kidney to body weight ratio at this timepoint compared to *Pkd1^+/+^* controls (**Fig.S3A**); an elevation which persisted up to 12 months of age (**Fig.S3B**). However, measuring BUN at intervals over 12 months revealed that this surrogate marker of renal function only increases from 12 months of age in *Pkd1^RC/RC^* mice (mean difference = 14.8mg/dL, 95% CI = 1.4-28.1, *p =* 0.023), with no significant change in BUN in the intervals measured in this murine cohort before 9 months (**Fig.3B**). Using the 488 nm laser of our lightsheet microscopy to image autofluorescence and delineate tubular epithelial microstructure, we were able to clearly visualise microscopic cysts in 3-month-old *Pkd1^RC/RC^* kidney vibratome slices, enabling quantification of cystic burden (**Fig.3C**). Across a total of 166 cysts measured across kidney slices from *n* = 3 *Pkd1^RC/RC^* mice, the mean cyst volume was calculated as 3.85 × 10^−5^ ± 2.93 × 10^−5^ mm^3^, and we found 2.32 ± 1.89% of the kidney volume was occupied by cysts in *Pkd1^RC/RC^* at this early timepoint.

**Fig. 3.**
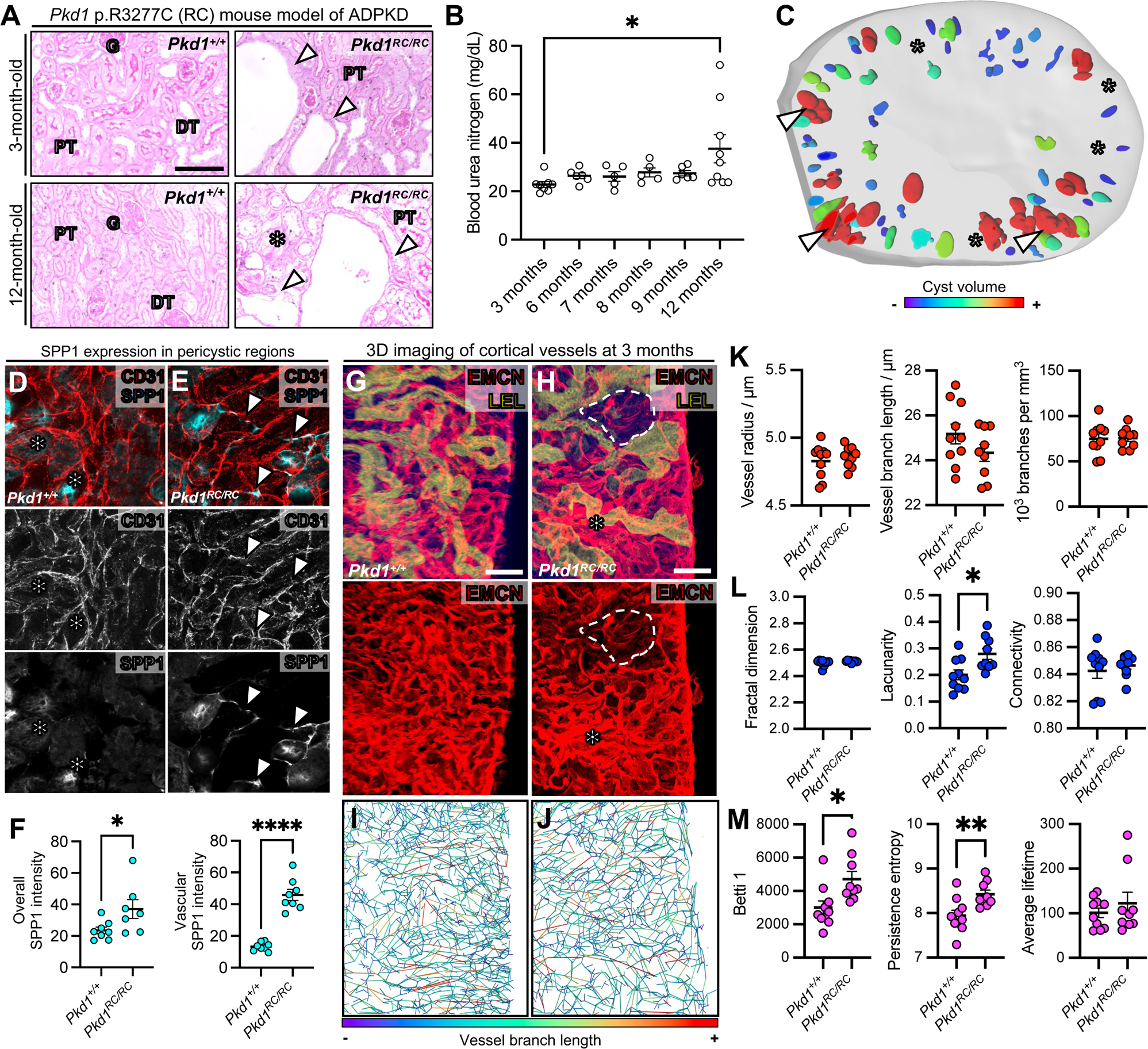
Identification of early structural and molecular changes to the vasculature in a mouse model of ADPKD. **(A)** Histology of the *Pkd1* p.R3277C (RC) mouse model of ADPKD shown by Periodic Acid-Schiff staining. Staining was performed at 3 months and 12 months, and each timepoint utilizing wildtype (*Pkd1^+/+^*) or homozygous (*Pkd1^RC/RC^*) littermates. Representative of four slides per kidney, with *n =* 4 mice analysed per group. Scale bar = 100 μm. **(B)** Calculation of blood urea nitrogen (BUN) across a timeseries in *Pkd1^RC/RC^* mice. Each point on the graph represents an individual mouse. ANOVA was used to compare groups and demonstrated a statistically significant difference (*F =* 2.6, *p =* 0.049). Upon Tukey’s multiple comparisons, only comparison of the 3-month and 12-month timepoint demonstrated statistical significance (mean difference = 14.8mg/dL, 95% CI = 1.4-28.1, *p =* 0.023). **(C)** 3D rendering of lightsheet images of cysts (coloured) within an optically cleared vibratome slice of a *Pkd1^RC/RC^* kidney at 3 months. The kidney tissue is shown in grey. Cysts are predominantly located in the cortex and their size is represented by a colourmap, with red being the greatest relative volume and purple being the smallest. Arrowheads represent cysts whereas asterisks represent regions adjacent to cysts. **(D)** Representative confocal microscopy of four images each taken from *n* = 3 *Pkd1^+/+^* mouse kidneys at 3 months of age stained for EMCN and SPP1 before optical clearing. The asterisks represent nonvascular SPP1-expression. **(E)** Representative confocal microscopy of four images each taken from *n* = 3 *Pkd1^RC/RC^* mouse kidneys at 3 months of age stained for EMCN and SPP1 before optical clearing. The white arrowheads indicate SPP1-expressing vasculature. **(F)** Quantification of overall SPP1 fluorescence intensity in images from (**D**) and (**E**). On the left, each point represents a region of interest imaged across the tissue, with *n* = 3 per group. On the right, individual vessel branches were assessed and pooled, such that each point represents an average fluorescence intensity of 10 vessel branches. Using Student’s *t* test, there was a statistically significant difference in both the overall SPP1 expression (mean difference 13.6 ± 6.0, 95% CI = 0.66-26.5, *p =* 0.041) and the vascular SPP1 expression (mean difference 33.6 ± 3.6, 95% CI = 24.8-40.3, *p <* 0.0001) between conditions. *: *p <* 0.0332, **: *p* < 0.0021, ***: *p* < 0.0002, ****: *p* < 0.0001. **(G)** 3D reconstruction of confocal images of the endomucin (EMCN)^+^ vasculature in *Pkd1^+/+^* kidney vibratome slices at 3 months of age. The kidneys were also stained with Tomato lectin (LEL) to discriminate tubules and orientate cortex from medulla. Representative of *n* = 4 kidneys imaged, with a minimum of 3 regions scanned per kidney. Scale bar = 50 μm. **(H)** 3D reconstruction of confocal images of the endomucin (EMCN)^+^ vasculature in *Pkd1^RC/RC^* kidney vibratome slices at 3 months of age. The kidneys were also stained with Tomato lectin (LEL) to discriminate tubules and orientate cortex from medulla. Representative of *n* = 4 kidneys imaged, with a minimum of 3 regions scanned per kidney. Scale bar = 50 μm. **(I)** VesselVio software was used to quantify vessel geometry of EMCN^+^ vasculature in the cortex of *Pkd1^+/+^* kidney vibratome slices at 3 months of age. This analysis was performed on all samples. Visual demonstration of the wire skeleton generated and colour-coded for vessel branch length. **(J)** VesselVio software was used to quantify vessel geometry of EMCN^+^ vasculature in the cortex of *Pkd1^RC/RC^* kidney vibratome slices at 3 months of age. This analysis was performed on all samples. Visual demonstration of the wire skeleton generated and colour-coded for vessel branch length. **(K)** Analysis of geometric quantities, including vessel radius, length, and density (defined as 10^3^ vessel branches per mm^3^ of tissue). Each point represents an individual ROI (minimum of *n =* 3 ROIs per mouse) imaged and pooled across *n* = 4 mice per group. Student’s *t* test demonstrated no significant differences between *Pkd1^+/+^* and *Pkd1^RC/RC^* kidneys at 3 months of age. **(L)** 3D fractal analysis including fractal dimension, lacunarity and connectivity, between *Pkd1^+/+^* and *Pkd1^RC/RC^* kidneys at 3 months of age. Each point represents an individual region of interest imaged and pooled across *n* = 4 mice per group. Student’s *t* test demonstrated that only lacunarity was significantly different between the two groups (mean difference = 0.079 ± 0.028, 95% CI = 0.020-0.14, *p =* 0.012). **(M)** Topological analysis including Betti 1, persistence entropy and average lifetime, between *Pkd1^+/+^*and *Pkd1^RC/RC^* kidneys at 3 months of age. Each point represents an individual region of interest imaged and pooled across *n* = 4 mice per group. Student’s *t* test demonstrated that both Betti 1 (mean difference = 1711 ± 604.3, 95% CI = 435.9-604.3, *p =* 0.012) and persistence entropy (mean difference = 0.48 ± 0.15, 95% CI = 0.16-0.81, *p =* 0.006) were significantly different between the two groups.

The above characterisation implies that 3 months is an early timepoint in cystic burden and disease severity of this model. We thus used 3D confocal microscopy to determine whether the unique subset of pericystic endothelium, characterised by SPP1 expression in end-stage human ADPKD kidneys, were also present in the *Pkd1^RC/RC^* mouse model. 500 μm thick vibratome sections were obtained, including cortex and medulla of *Pkd1^+/+^* or *Pkd1^RC/RC^* mouse kidneys at 3 months of age. 3D imaging of fixed *Pkd1^+/+^* or *Pkd1^RC/RC^*mouse kidney 500 μm vibratome sections was performed at 3 months of age, after staining for CD31 and SPP1 and optical clearing. Considering that ∼98% of the kidney volume was non-cystic, we used the 488 nm laser for orientation within the kidney and visualisation of cysts. In this way, we were able to focus on regions adjacent to cysts within the kidney, considering that these regions in human ADPKD kidneys contained SPP1^+^ endothelia. Scant SPP1 expression was detected in CD31^+^ capillary vasculature within *Pkd1^+/+^*kidneys at 3 months (**Fig.3D**). Conversely, SPP1^+^ ECs were found interspersed amongst the peritubular capillary network in 3-month-old *Pkd1^RC/RC^* mouse kidneys (**Fig.3E**). The fluorescence intensity of SPP1 was quantified from these 3D images, both across the kidney volume and individually within CD31^+^ vasculature (**Fig.3F**). When averaged across the kidney volume, the SPP1 fluorescence intensity subtly, but significantly increased in *Pkd1^RC/RC^* mouse kidneys as compared with wildtype controls (mean difference 13.6 ± 6.0, 95% CI = 0.66-26.5, *p =* 0.041). The mean SPP1 fluorescence intensity of CD31^+^ vasculature, however, increased over three-fold in homozygous mice compared to wildtype controls (mean difference 33.6 ± 3.6, 95% CI = 24.8-40.3, *p <* 0.0001). Thus, akin to the human disease, endothelium expressing SPP1 os present within the pericystic microenvironment in an orthologous mouse model of ADPKD, but at early stages long before the decline of renal function in this model.

We then characterised if the presence of pericystic endothelium at an early disease stage in *Pkd1^RC/RC^* mice was associated with structural abnormalities of the renal microvasculature. To do this, we immunostained for the blood microvascular marker, EMCN, whilst simultaneously labelling these tissues with Tomato lectin, before high-resolution 3D imaging deep into the tissue using a confocal microscope (48). In our hands, wholemount labelling using Tomato lectin labelled tubular epithelia, which enabled clear demarcation of the cortex and medulla using 3D microscopy (**Fig.S4A**), with cysts demarcated by autofluorescence (**Fig.S4B**). Maximum intensity projection of the EMCN^+^ vasculature in the cortex demonstrated regularly patterned peritubular capillaries in *Pkd1^+/+^* kidneys (**Fig.3G**). Visually, there was non-uniformity in the patterning of the vasculature in the *Pkd1^RC/RC^* kidney cortex at three months. This non-uniformity was pronounced in the subregions adjacent to cysts, which contained an abundance of disorganised EMCN^+^ capillaries (**Fig.3H**).

Given the visually abnormal patterning corresponding to pericystic endothelium, we analysed renal blood microvascular geometry by segmenting the intact 3D network, before using open source 3D vascular analysis software (56) to quantitate conventional vessel branch geometric properties including the branch length (mapped and visualised using 3D rendering in **Fig.3I-J**), radius and vascular density; the number of branches per mm^3^ of tissue. There were no significant differences in EMCN^+^ microvessels branch radius, length or density between *Pkd1^+/+^* or *Pkd1^RC/RC^* mouse kidneys at 3 months (**Fig.3K**). A limitation of such analyses is that they do not consider the vascular network patterning, and thus may not capture visually detectable phenotypes. We therefore turned to a novel approach of analysing the kidney microvasculature in the ADPKD mouse model. We leveraged 3D fractal analysis, which better represents the patterning and complexity of the network in 3D space, as compared with individual summary statistics such as mean radius and length. Using 3D imaging of EMCN^+^ microvessels, we were able to quantify metrics such as fractal dimension; representing vascular complexity, lacunarity; representing non-uniformity, and connectivity. This analysis demonstrated a significant increase in the heterogeneity of distribution of the vascular network in *Pkd1^RC/RC^*mouse kidneys as compared with wildtype controls at 3 months (mean difference in lacunarity = 0.079 ± 0.028, 95% CI = 0.020-0.14, *p =* 0.012), without significant changes in complexity (fractal dimension) or connectivity between conditions (**Fig.3L**), agreeing with the visual changes observed in 3D microscopy of a more non-uniform vascular network in mutant mice, which may have implications for blood perfusion within the cystic kidney.

To obtain deeper insights into the nature of vascular patterning in the ADPKD mouse model, we derived topological measures from segmented images of the EMCN^+^ microvasculature, including Betti numbers; representing the number of vascular loops, persistence entropy; representing vascular disorganisation, and average lifetime; representing persistence of patterns in 3D space. We found both number of loops (mean difference of Betti 1 = 1711 ± 604.3, 95% CI = 435.9-604.3, *p =* 0.012) and vascular disorganisation (mean difference of persistence entropy = 0.48 ± 0.15, 95% CI = 0.16-0.81, *p =* 0.006) to be significantly greater in *Pkd1^RC/RC^*mouse kidneys as compared to wildtype controls at 3 months, with no significant change in the persistence of patterns in 3D space (average lifetime, **Fig.3M**). By contrast, our 3D analysis of geometric (**Fig.S5A**), fractal (**Fig.S5B**) or topological (**Fig.S5C**) measures detected no significant changes in the EMCN^+^ medullary vasculature of *Pkd1^+/+^* or *Pkd1^RC/RC^* mouse kidneys.

Altogether, we have shown that pericystic endothelium expressing SPP1, present in end-stage ADPKD in humans, is also identifiable from an early stage in *Pkd1^RC/RC^* mouse kidneys, preceding decline in renal function. By applying novel 3D analysis of the kidney microvasculature, we demonstrate that pericystic regions in ADPKD; mapping to SPP1^+^ ECs, are associated with heterogeneous patterning, with disorganisation and a greater number of capillary loops; altogether indicating aberrant kidney microvascular remodelling as a feature of early ADPKD.

### Early reduction in cortical blood flow independent of cysts in murine PKD

Having identified the unique molecular signature of an endothelial cell subset in ADPKD, with its identification associated with aberrant structural remodelling early in the *Pkd1^RC/RC^* mouse model, we sought to assess whether blood perfusion within the kidney is concurrently altered. To do this, we leveraged a 9.4T MRI scanner to perform *in vivo* imaging of live *Pkd1^+/+^* or *Pkd1^RC/RC^* mice (**Fig.4A**). We applied a multimodal contrast-free approach, firstly assessing the structure of the kidney using a T2-FLAIR sequence, before using ASL to map and quantify RBF (82). As T2-FLAIR was able to discriminate cortex and medulla within the kidney, aligned ASL images from the same mice could then be used to discriminate blood flow originating from cortex or medulla. *Pkd1^+/+^* or *Pkd1^RC/RC^*mice were scanned at 9 months of age; a timepoint which precedes the significant increase in BUN we observed in this model. We also scanned mice at the earlier timepoint of 3 months, corresponding to the detection of pericystic microvessels and aberrant vascular patterning within the ADPKD microenvironment.

**Fig. 4.**
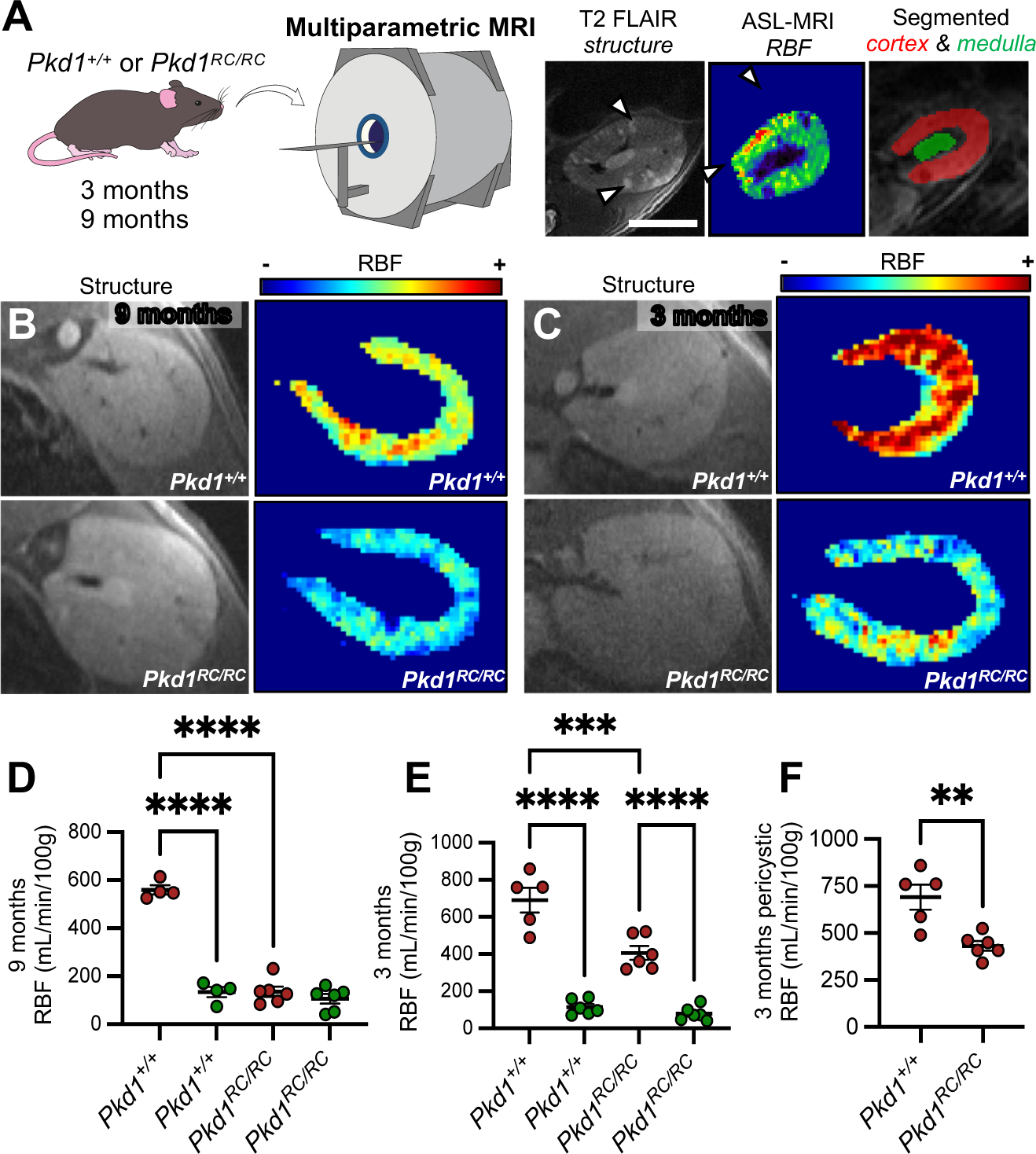
Multiparametric MRI shows early and regional reduction in renal blood flow in mice with ADPKD. **(A)** Schematic showing the experimental setup, with *Pkd1^+/+^* or *Pkd1^RC/RC^* mice used for T2 FLAIR magnetic resonance imaging (MRI) to capture renal structure before arterial spin labelling (ASL)-MRI was used to assess renal blood flow (RBF). Alignment of imaging facilitated differentiation between cortical and medullary RBF values. Scale bar in T2 FLAIR image = 2 mm. **(B)** Representative ASL-MRI imaging at 9 months of age in *Pkd1^+/+^* (*n* = 4) and *Pkd1^RC/RC^* mice (*n* = 6). Heatmapping RBF values visually demonstrates local reduction in flow within the cortex of *Pkd1^RC/RC^* mouse kidneys. **(C)** Representative ASL-MRI imaging at 3 months of age in *Pkd1^+/+^* (*n* = 5) and *Pkd1^RC/RC^* mice (*n* = 6). Heatmapping RBF values visually demonstrates local reduction in flow within the cortex of *Pkd1^RC/RC^* mouse kidneys. **(D)** Comparison of RBF values in cortex (red points) and medulla (green points) at 9 months of age across conditions. Each point represents a single mouse scanned. ANOVA demonstrated a significant difference across conditions, with a difference between the cortex and medulla of wildtype mice (mean difference = 426.1 ± 31.1 mL/min/100g, 95% CI = 331.4-529.8, *p <* 0.0001) as well as a significant reduction in the cortex of *Pkd1^RC/RC^* as compared to *Pkd1^+/+^* mouse kidneys (mean difference = 424.4 ± 30.2 mL/min/100g, 95% CI = 337.9-510.0, *p <* 0.0001). **(E)** Comparison of RBF values in cortex (red points) and medulla (green points) at 3 months of age across conditions. Each point represents a single mouse scanned. ANOVA demonstrated a significant difference across conditions, with a difference between the cortex and medulla of wildtype mice (mean difference = 577.6 ± 53.0 mL/min/100g, 95% CI = 428.6-726.6, *p <* 0.0001) as well as a significant reduction in the cortex of *Pkd1^RC/RC^* as compared to *Pkd1^+/+^* mouse kidneys (mean difference = 285.0 ± 52.9 mL/min/100g, 95% CI = 136.1-434.0, *p =* 0.0002). **(F)** Comparison of RBF values in cortex at 3 months of age across conditions, after segmentation and analysis of pericystic regions of the cortex devoid of cysts in the mutant mice. Each point represents a single mouse scanned. Student’s *t* test demonstrated a significant reduction in RBF in *Pkd1^RC/RC^* as compared to *Pkd1^+/+^* mouse kidneys (mean difference in 259.6 ± 66.4 mL/min/100g, 95% CI = 109.5-409.7, *p =* 0.0035). **: *p* < 0.0021, ***: *p* < 0.0002, ****: *p* < 0.0001.

When visualised using heatmaps, there were striking changes to RBF within the cortex of *Pkd1^RC/RC^* mice as compared to wildtypes at both 9 months (**Fig.4B**) and 3 months (**Fig.4C**). We then discerned average values of RBF from each ASL image, analysing cortex and medulla of each kidney separately. There was significantly higher RBF in the cortex as compared to the medulla (**Fig.4D**) at both 9 months (mean difference = 426.1 ± 31.1 mL/min/100g, 95% CI = 331.4-529.8, *p <* 0.0001) and 3 months (mean difference = 577.6 ± 53.0 mL/min/100g, 95% CI = 428.6-726.6, *p <* 0.0001) in *Pkd1^+/+^*wildtype mice, RBF values which corresponded to those reported in the literature (44, 83, 84). We then compared between health and disease, revealing a 76% reduction in mean cortical blood flow at 9 months in *Pkd1^RC/RC^* mice (**Fig.4D**, mean difference = 424.4 ± 30.2 mL/min/100g, 95% CI = 337.9-510.0, *p <* 0.0001). Strikingly this significant reduction was also observed at the earlier timepoint of 3 months, with a 35% reduction in mean cortical RBF in cystic mice (**Fig.4E**, mean difference = 285.0 ± 52.9 mL/min/100g, 95% CI = 136.1-434.0, *p =* 0.0002). Medullary blood flow, however, was not significantly different between *Pkd1^RC/RC^* or *Pkd1^+/+^* mice at either timepoint.

The local reduction in blood flow within the cortex of *Pkd1^RC/RC^* mouse kidneys corresponds to the localisation of pericystic endothelium and aberrant vascular remodelling within the microenvironment adjacent to cysts. We thus leveraged the paired T2-FLAIR structural imaging to segment regions adjacent to macroscopic cysts within *Pkd1^RC/RC^*kidneys, and quantified blood flow within these regions, comparing this to cortical subregions of *Pkd1^+/+^* controls (**Fig.4F**). We found that the significant reduction of RBF observed between *Pkd1^RC/RC^*and *Pkd1^+/+^* kidneys was maintained using this approach, with pericystic subregions of *Pkd1^RC/RC^* mouse cortex having a 38% reduction on average compared to control cortex (mean difference in 259.6 ± 66.4 mL/min/100g, 95% CI = 109.5-409.7, *p =* 0.0035). Thus, the reduction in RBF in the cortex of *Pkd1^RC/RC^* mouse kidney, occurs in regions adjacent to cysts, with associated molecular and structural vascular changes at an early stage of the disease preceding decline in renal function.

## DISCUSSION

This work leveraged a range of parallel approaches, including transcriptomics at cellular resolution, 3D imaging of intact, immunolabelled and kidney tissues, and *in vivo* imaging of kidney structure and renal blood flow. Collectively, our data show that the kidney blood vasculature is perturbed at molecular, structural and functional levels from the early stages of ADPKD. We detected blood microvessels in the pericystic regions of patient’s kidneys; with a molecular phenotype which is not observed in chronic kidney disease or transplant rejection. This molecular phenotype, suggesting abnormal metabolism and angiogenesis, is distinct from the common transcriptional signature of endothelial injury observed in kidney disease. By identifying SPP1 as a marker of this pericystic endothelium at transcriptome and protein levels, we detected its presence from as early as 3 months in an orthologous murine model of ADPKD. This phenotype occurred concurrent with aberrant structural remodelling identified by 3D quantitative analysis of kidney microvasculature using fractal and topological approaches. The aberrant endothelial phenotype mapped to the microenvironment surrounding cysts, where there is also local impairment of blood flow in *Pkd1^RC/RC^*mouse kidneys. Critically, these changes occur prior to the onset of overt decline in kidney function in mice, implicating alterations to the kidney vasculature as a potential therapeutic target in ADPKD at stages where kidney function is still intact.

This study, bridging molecular, structural, and functional levels, sheds new lights on the phenotype of the kidney vasculature in ADPKD. Although it has been established that mutations in *Pkd1* cause defects in the development (85–87) and flow-mediated dilation of the systemic vasculature (23), the phenotype of the renal microvasculature in ADPKD has remained somewhat elusive. To our knowledge, these experiments are the first to interrogate the molecular phenotype of the vasculature of the human cystic kidney in depth. By generating and validating a reference scRNA-seq atlas of the kidney microvasculature, applicable to other kidney diseases, we dissected endothelial subtype-specific transcriptomes from human ADPKD snRNA-seq data (34). In so doing, we identified a unique cluster of endothelium expressing SPP1 present in human ADPKD, and within the *Pkd1^RC/RC^*mouse model at the 3-month timepoint. One could postulate that the lack of such a molecular phenotype in normal kidney, or other kidney pathologies in our scRNA-seq atlas, could potentially be explained by the intrinsic effect that PKD1 mutation or deletion has on endothelial development and function (23, 24, 29). Our 3D interrogation of vascular architecture within pericystic regions at this early stage of disease revealed disorganisation, with an increased number of loops, manifesting in heterogenous patterning. Our findings reconcile previous angiography, immunohistochemistry and corrosion casting of human end-stage ADPKD explants and mouse models – with some reports indicating angiogenic remodelling as a feature of ADPKD (25, 29, 88), whereas others suggest a lower overall renal vascular density in cystic disease (25, 27, 28). We found that conventional, 3D geometric analysis of the microvasculature failed to find any changes in metrics such as branch radius, length, or density, which is why we developed a more sensitive 3D approach to detect overall patterning metrics of the kidney microvasculature, incorporating topological information and fractal analysis. The aberrant and structurally malformed pericystic microvascular subset could represent a compensatory response to the inflammatory microenvironment within the pericystic regions of ADPKD kidneys (16). This response may be impaired, as evidenced by reduction in VEGFR expression in pericystic endothelium, a finding also reflected by previous work showing loss of VEGF signalling as a feature of PKD rodent models (29, 89). Alternatively pericystic endothelium, in its altered phenotypic state, could contribute to further inflammation and fibrotic remodelling, by driving the recruitment and activation of adjacent cells, including fibroblasts (17) or macrophages (90, 91). In any case, these vessels are associated with a decline in local cortical perfusion within the kidney, as we demonstrated using ASL of *Pkd1^RC/RC^* mouse kidneys at the early, 3-month stage of the disease.

This study generated technical advances, such our scRNA-seq atlas, which alongside new studies (92), provides a molecular reference for future transcriptomic experiments interrogating the kidney vasculature in health and disease. The novel 3D approach to quantitatively assess topological and fractal information from the kidney vasculature has implications in studying the vasculature in not only the kidney, but other situations featuring vascular remodelling. The present experiments also have clinical implications. In our study, the discovery of pericystic endothelium, with associated structural and vascular functional defects, has been performed both in humans and at the early stages of a clinically relevant, orthologous mouse model, at a timepoint where ∼98% of the kidney volume remains unoccupied by cysts; sufficient normal kidney parenchyma to maintain baseline blood urea nitrogen values. The temporal progression of disease in *Pkd1^RC/RC^* mice is comparable with human ADPKD (32), and hints towards the presence of renal microvascular abnormalities at molecular, structural and functional levels prior to inexorable decline in kidney function of patients. Indeed, renal perfusion has been detected in patients with ADPKD using contrast-enhanced MR angiography, and flow detected within the renal arteries is an independent predictor of kidney function (93, 94). Our study strengthens the case for renal microvascular dysfunction being an early hallmark of ADPKD; using a non-contrast approach to extend these previous findings and demonstrating intrinsic microvascular dysfunction of the organ. The insufficient resolution of conventional *in vivo* imaging approaches used in patients makes it challenging to detect such regional defects in perfusion. It would be prudent to apply such techniques to future clinical trials of ADPKD, monitoring the response of intrinsic renal blood flow to pharmacological agents. To mitigate the limitations of MRI, such as partial volume effect due to averaging ‘zero’ flow regions of epithelial cysts within renal blood flow measurements, alternative and emerging techniques, such as photoacoustic imaging (27) or multiphoton microscopy (95), could be used to non-invasively assess kidney vasculature paired with novel 3D analysis approaches to assess vascular patterning described in this paper. Our findings also have therapeutic implications. It has already been established that manipulation of VEGF signalling alters disease phenotype in rapidly progressive rodent models of PKD (29, 89, 96). TIE-mediated angiopoietin signalling was revealed as another potential candidate to target aberrant vascular remodelling, which has been investigated in the context of other kidney diseases (97–100) but not ADPKD. Importantly, our findings unearth vascular heterogeneity in ADPKD which may be amenable to targeted approaches, such as protein carriers (101), nanoparticles (102) or gene therapy which could realise the potential of targeting the vasculature in ADPKD whilst reducing systemic unwanted side-effects.

### Conclusion

In summary, we resolve the molecular, structural, and functional phenotype of the renal microvasculature in ADPKD, manifesting in defective molecular, aberrantly remodelled, and poorly functional pericystic endothelium. The presence of the pericystic endothelial phenotype precedes inexorable decline in renal function. Our findings strengthen the case for the renal microvasculature as a therapeutic target in ADPKD within a timeframe during which renal function is still salvageable. Given the new evidence presented in this study, the time is drawing nearer for the development and implementation of vascular-based pharmacological approaches as a new therapeutic direction for ADPKD.

## AUTHOR CONTRIBUTIONS

DJ, PS, BH, ML, SWS and DAL conceived the study. Acquisition of human or mouse material, mouse husbandry, renal function assays and immunostaining or histology were performed by DJ, MB, LW, WM, EF, MKJ, RP, SM, KP, RM and JC. Confocal or lightsheet imaging and 3D analysis were performed by DJ, GP, BD, HM, RL, SN, CW and DM. Analysis of single-cell and single-nucleus transcriptomics data was performed by DJ, MB, GP, BS and YM. MRI experiments and analyses were performed by DJ, CP and DT. Project oversight and supervision was provided by AW, PW, PS, RH, MRC BH, ML, SWS and DAL. DJ wrote the first draft of the paper, refined by DAL, and subsequently all authors were involved in revision and preparation of the final manuscript for submission.

## ACKNOWLEDGEMENTS

DJ was supported by a Rosetrees Trust PhD Plus Award (PhD2020\100012), a Foulkes Foundation Postdoctoral Fellowship and the Specialised Foundation Programme in the East of England Foundation Schools. The work is also supported by project grants from the Medical Research Council (MR/P018629/1 and MR/J003638/1) an access grant at the UKRI’s Octopus Central Laser Facility (23130027), a PKD Charity Grant (PKD-22-01) and a Wellcome Trust Investigator Award (220895/Z/20/Z) to DAL. DAL’s laboratory is supported by the NIHR Biomedical Research Centre at Great Ormond Street Hospital for Children NHS Foundation Trust and University College London. We thank Peter Harris (Mayo Clinic) for Pkd1^RC^ mice.

## SUPPLEMENTARY DATA LEGENDS AND TABLES

**Fig.S1. Profiling marker gene expression of vascular endothelial subsets across the human kidney vascular single-cell transcriptome**

**(A)** Discrimination of endothelial cells in the UMAP of kidney single-cell atlas was performed by assessing cells expression CDH5 and FLT1, both blood endothelial cell markers.

**(B)** Feature plots showing expression of SSUH2, FBLN2 and GPM6A across the dataset, demonstrated restricted expression within the endothelial cluster.

**(C)** Dot plot showing consistent expression of above selected marker genes across the five individual datasets used to construct the human kidney vascular endothelial atlas.

**Fig.S2. Annotation and characterization of atlas of blood endothelial cells in human ADPKD**

**(A)** Individual samples used to construct the dataset, including 5 control kidneys and 8 kidney explants with ADPKD.

**(B)** Dot plots showing top 2-3 marker genes per cluster, with selected candidates *SSUH2*, *FBLN2* and *GPM6A* annotating corresponding cell types within this new dataset.

**(C)** Bar chart showing relative abundance of cell types across conditions. Injured endothelial cells and cystic endothelial cells are unique to ADPKD samples.

**(D)** Violin plot showing differential expression analysis of arterial, efferent, and afferent arteriolar, glomerular capillary and ascending vasa recta endothelial cells between conditions. Each point represents a gene, with red points being differentially expressed and statistically significant, blue not meeting criteria for significance and black not differentially expressed.

**(E)** Dot plot of DEGs identified in pericystic endothelium from the snRNA-seq ADPKD atlas, analyzing cell types within the prior scRNA-seq kidney endothelial cell reference atlas for expression of these transcripts.

**(F)** Violin plot of vascular tyrosine kinases expressed by pericystic endothelium, at lower levels on average, as compared to other EC types within the snRNA-seq ADPKD atlas. These include members of the vascular endothelial growth factor receptor (VEGFR) family and members of the tyrosine kinase with immunoglobulin like and EGF like domains (TIE) family.

**Fig.S3. Characterisation of kidney to bodyweight ratios in the mouse model of ADPKD**

**(A)** Assessment of kidney to bodyweight ratio at 3 months in *Pkd1^RC/RC^* as compared to *Pkd1^+/+^* mouse kidneys. Each point on the graph represents an individual mouse analysed. Student’s *t* test demonstrated a statistically significant increase in mutant as compared to wildtype mice (mean difference in 0.87 ± 0.13% kidney weight, 95% CI = 0.58-1.15, *p <* 0.0001).

**(B)** Assessment of kidney to bodyweight ratio at 12 months in *Pkd1^RC/RC^* as compared to *Pkd1^+/+^* mouse kidneys. Each point on the graph represents an individual mouse analysed. Student’s *t* test demonstrated a statistically significant increase in mutant as compared to wildtype mice (mean difference in 1.6 ± 0.53% kidney weight, 95% CI = 0.39-2.81, *p =* 0.016).

**Fig.S4. Three-dimensional confocal imaging of the kidney vasculature and cysts in cortex and medulla of an ADPKD mouse model**

**(A)** Representative maximum intensity projection of *n =* 4 kidneys from *Pkd1^+/+^* mice. Kidneys were stained for endomucin (EMCN), tomato lectin (LEL) and we also imaged autofluorescence to capture tissue architecture. The dotted line separated cortex from medulla, clearly demonstrated by the pattern of LEL staining.

**(B)** Representative maximum intensity projection of *n =* 4 kidneys from *Pkd1^RC/RC^* mice. Kidneys were stained for EMCN), LEL and we also imaged autofluorescence to capture tissue architecture. The dotted line separated cortex from medulla, clearly demonstrated by the pattern of LEL staining. Arrowhead show cysts within the autofluorescence channel. Scale bar = 1 mm.

**Fig.S5. Quantitative analysis of the kidney vasculature demonstrates no differences in the medulla of a mouse model of ADPKD.**

**(A)** Analysis of geometric quantities, including vessel radius, length and density (defined as 10^3^ vessel branches per mm^3^ of tissue). Each point represents an individual region of interest imaged within the medulla and pooled across *n* = 4 mice per group. Student’s *t* test demonstrated no significant differences between *Pkd1^+/+^* and *Pkd1^RC/RC^* kidneys at 3 months of age.

**(B)** 3D fractal analysis including fractal dimension, lacunarity and connectivity, between *Pkd1^+/+^* and *Pkd1^RC/RC^* kidneys at 3 months of age. Each point represents an individual region of interest imaged within the medulla and pooled across *n* = 4 mice per group. Student’s *t* test demonstrated no significant differences between *Pkd1^+/+^* and *Pkd1^RC/RC^* kidneys at 3 months of age.

**(C)** Topological analysis including Betti 1, persistence entropy and average lifetime, between *Pkd1^+/+^*and *Pkd1^RC/RC^* kidneys at 3 months of age. Each point represents an individual region of interest imaged within the medulla and pooled across *n* = 4 mice per group. Student’s *t* test demonstrated no significant differences between *Pkd1^+/+^* and *Pkd1^RC/RC^* kidneys at 3 months of age.

**Table S1. List of differentially expressed genes within the scRNA-seq atlas of kidney endothelial cells in normal and diseased kidneys**

Differential expression analysis was performed using the Wilcoxon rank sum test. Tables are presented with gene names, *p* value (p_val), log_2_FC (avg_log2FC), percentage of cells within the cluster expressing the gene of interest (pct.1), percentage of cells across all other clusters expressing the gene of interest (pct.2), adjusted *p* value (p_val_adj) and name of cell type (cluster).

**Table S2. List of differentially expressed genes within the snRNA-seq atlas of kidney endothelial cells in ADPKD**

Differential expression analysis was performed using the Wilcoxon rank sum test. Tables are presented with gene names, *p* value (p_val), log_2_FC (avg_log2FC), percentage of cells within the cluster expressing the gene of interest (pct.1), percentage of cells across all other clusters expressing the gene of interest (pct.2), adjusted *p* value (p_val_adj) and name of cell type (cluster).

**Table S2. List of genes differentially expressed by pericystic endothelium within the snRNA-seq atlas of kidney endothelial cells in ADPKD**

Differential expression analysis was performed using the Wilcoxon rank sum test. Tables are presented with gene names, *p* value (p_val), log_2_FC (avg_log2FC), percentage of cells within the cluster expressing the gene of interest (pct.1), percentage of cells across all other clusters expressing the gene of interest (pct.2), adjusted *p* value (p_val_adj) and name of cell type (cluster). The first tab in the Excel file contains genes which are expressed, on average, at higher levels compared to other EC types. The second tab contains genes which are expressed, on average, at lower levels compared to other EC types.

