## Supplementary figures and images for "A unique subset of pericystic endothelium associates with aberrant microvascular remodelling and impaired blood perfusion early in polycystic kidney disease"

### Supplementary Figure 1

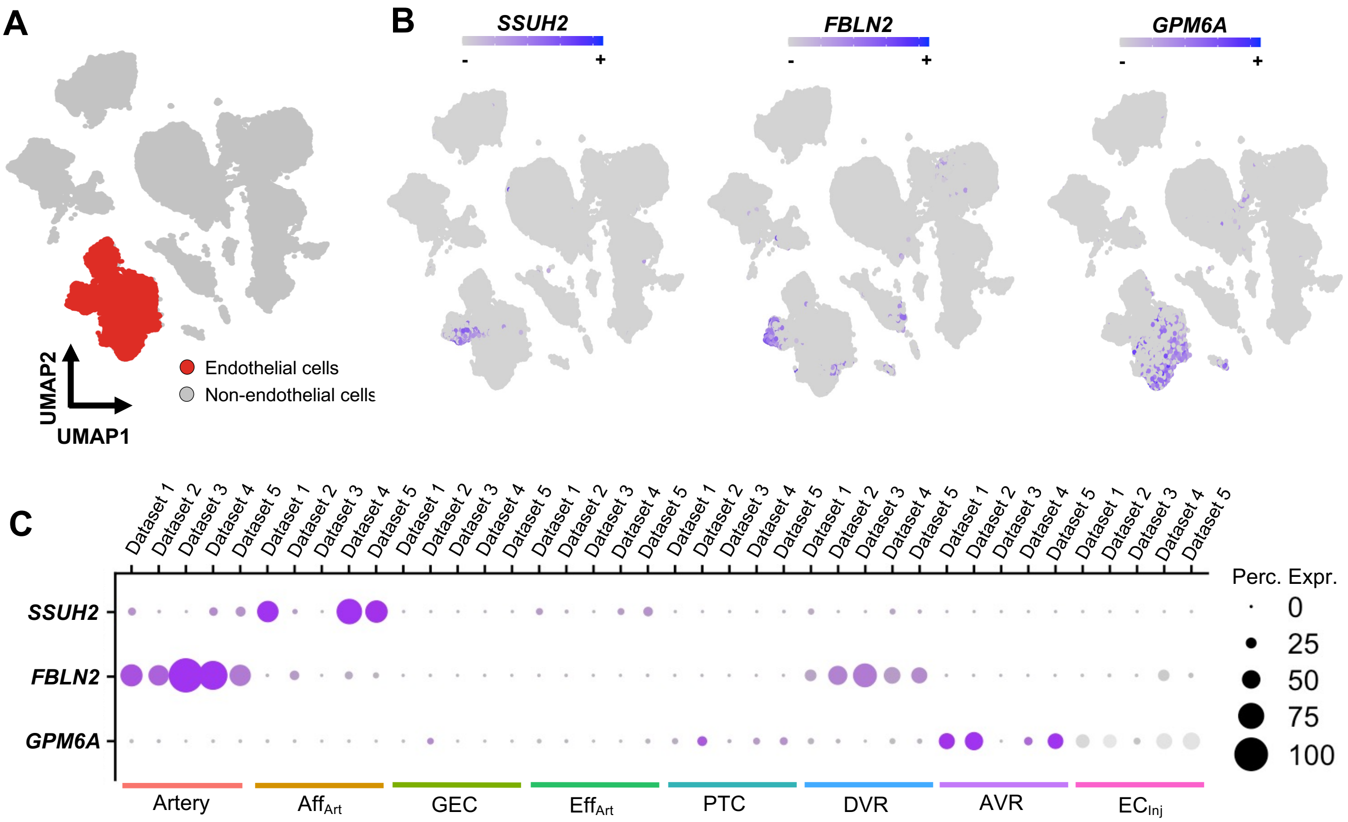

### Supplementary Figure 2

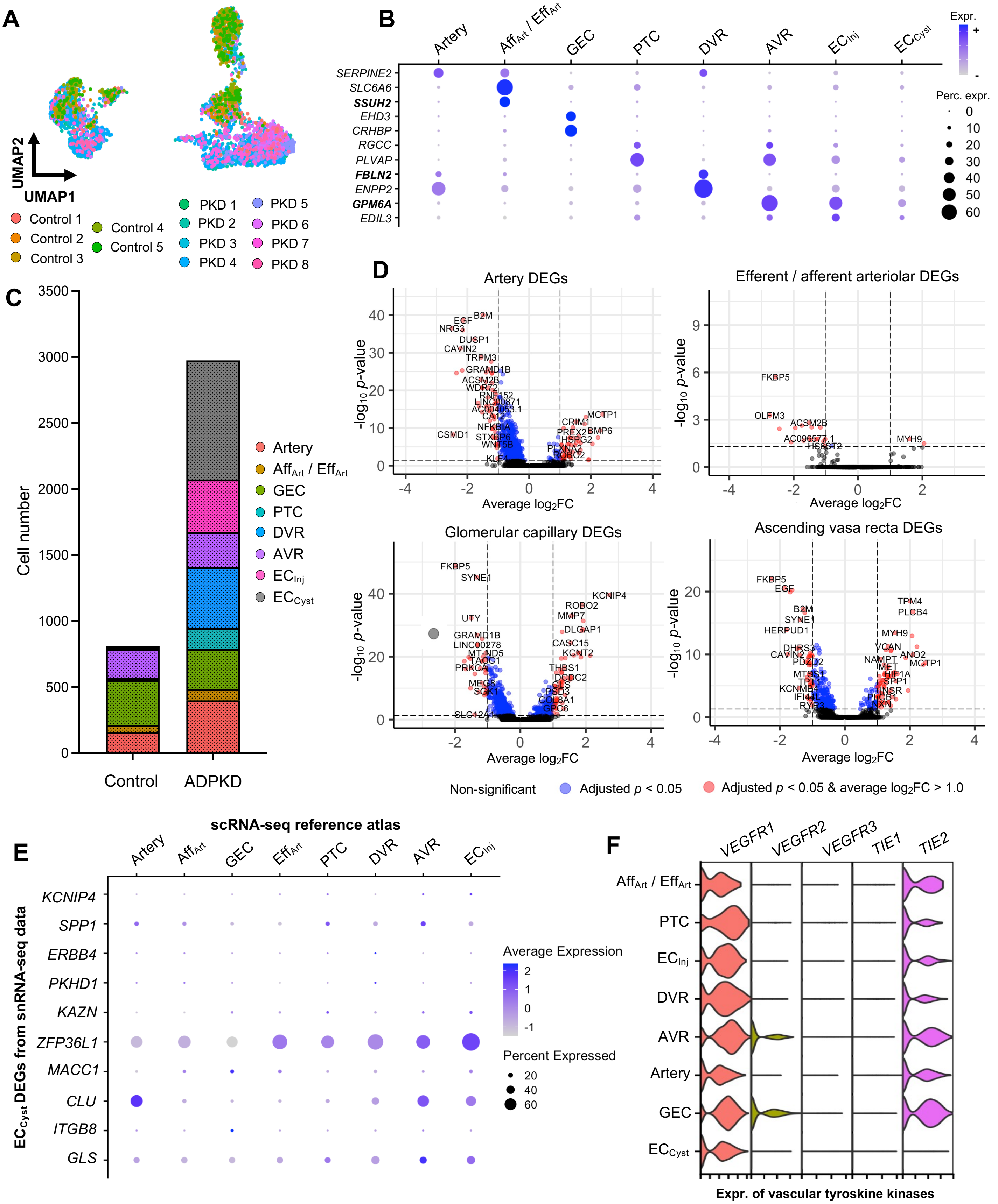

### Supplementary Figure 3

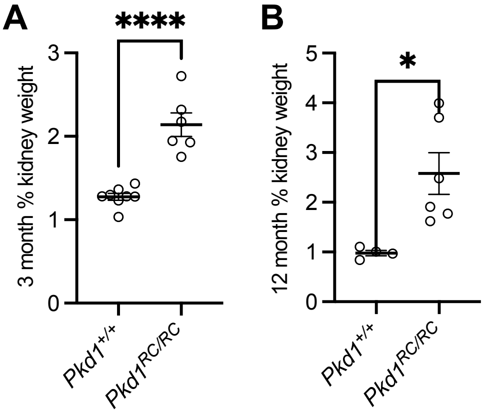

### Supplementary Figure 4

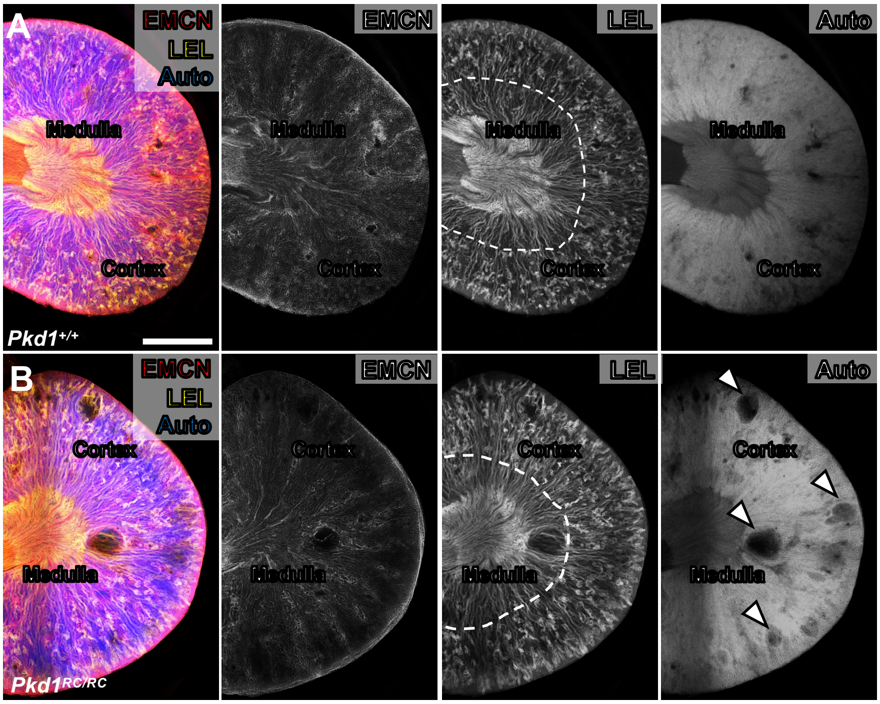

### Supplementary Figure 5

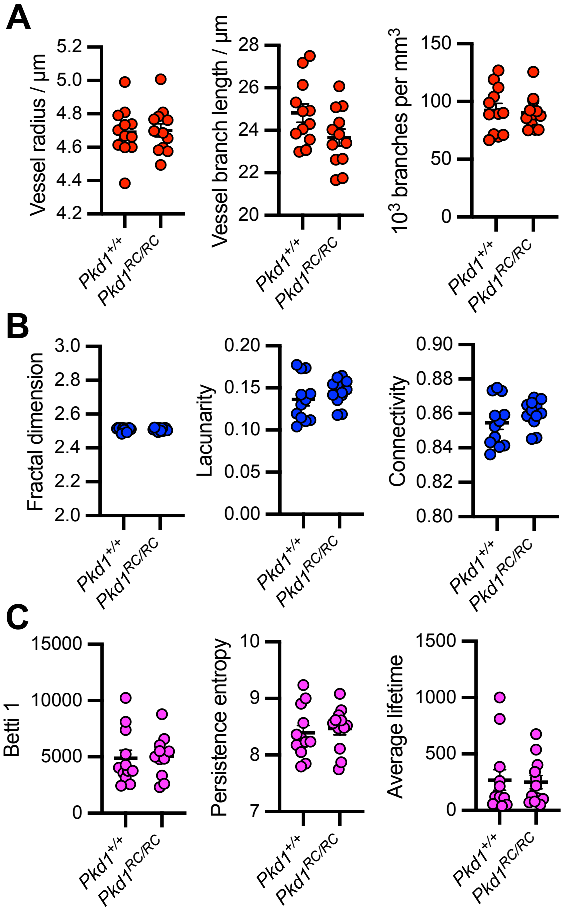
